# Ultraviolet B acts as a dietary restriction mimetic by targeting mitochondrial bioenergetics

**DOI:** 10.1101/2024.03.05.583543

**Authors:** Asya Martirosyan, Yuting Li, Yvonne Woitzat, Seunghye Lee, Li Fu, Maria A. Ermolaeva

**Affiliations:** Leibniz Institute on Aging – Fritz Lipmann Institute (FLI), Beutenbergstrasse 11, 07745, Jena, Germany; Shenzhen University Cancer Research Centre, Shenzhen University Xili Campus, 1066 Xueyuan Ave, Nanshan District, Shenzhen, China; Cluster of Excellence Cellular Stress Responses in Aging-Associated Diseases, University of Cologne, Joseph-Stelzmann-Straße 26, 50931 Cologne, Germany

## Abstract

Ultraviolet (UV) light is a common environmental stimulus, and UV exposure confers health benefits, with cellular targets still unclear. Here, we show that ultraviolet B (UVB) exposure alters mitochondrial bioenergetics in *C. elegans* and human skin fibroblasts triggering loss of membrane potential, mitochondrial fission and calcium release. This initial stress is followed by a recovery process relying on mitochondrial biogenesis and fusion, which prevents lasting mitochondrial damage. Strikingly, the transient decline of ATP synthesis caused by UVB-induced mitochondrial changes triggers a swift metabolic re-wiring response that resembles effects of dietary restriction (DR) at the organismal and molecular levels. Both recovery from UVB and DR-mimetic UVB effects require mitochondrial fusion, and we found that dysfunction of fusion during aging abrogates UVB benefits and sensitizes old nematodes to UVB toxicity. Finally, UVB irradiation of the skin was effective in inducing organismal fasting-like phenomena in proof-of-concept tests in young mice. We thus uncovered a novel evolutionary conserved cellular mechanism connecting UV light and metabolism. Our findings illuminate potential DR-mimetic properties of UVB and explain late life-specific UVB intolerance.

## Main text

Ultraviolet (UV) light is a ubiquitous environmental stimulus with substantial impact on human health ranging from skin carcinogenesis^1, 2^ to broad homeostatic benefits^3^. The studies linking UV and organismal homeostasis have mainly focused on its role in vitamin D synthesis, which cannot, however, explain the full spectrum of cellular and organismal changes observed upon UV exposure^4–6^. In previous work, we used *C. elegans* to uncover that transient DNA damage induced by ultraviolet-B (UVB) and ionizing radiation (IR) in actively proliferating cells leads to systemic stress tolerance via conserved MAP kinase signaling and activation of the ubiquitin proteasome system^7^. The contribution of mitochondria to this process was however unexplored in the previous study. Mitochondria is a potent signaling hub in the cell, which integrates various stress and metabolic challenges and instigates adaptive responses via mechanisms such as metabolic remodeling, mitophagy and mitochondrial unfolded protein response (UPR MT)^8, 9^. On the one hand, mitochondria possess their own genome that can be directly impacted by genotoxic treatments like UVB and IR^10^ and, additionally, nuclear DNA damage is known to interfere with cellular energy homeostasis and mitochondrial stress responses^11, 12^. In this study, we therefore decided to test the role of mitochondrial signaling in the adaptive stress responses to UVB and IR.

### Mitochondrial fusion mediates homeostatic recovery upon ultraviolet B exposure

Because mitochondrial fission and fusion play a key role in integrating mitochondrial stress responses with cellular energy states^13^, we decided to test if inhibition of either of these processes would have an impact on UVB- and IR-induced stress tolerance of *C. elegans*. Initially, we exposed L4 larval stage animals to 90Gy IR and different doses of UVB and 48h later treated them with heat stress, as described previously^7^, to identify the optimal UVB treatment conditions for the induction of adaptive stress tolerance by the specific device we used (Figure S1A). We next successfully confirmed the ability of the chosen UVB and IR doses to protect nematodes from stress by complementing heat stress tests with exposure to additional cellular stressors such as oxidative stress (induced by paraquat)^7^ and protein folding stress in the endoplasmic reticulum (ER) (induced by DTT)^14^ (Figure 1A and 1B). Subsequently, we asked if inhibition of mitochondrial fission by loss of the *C. elegans* DNM1L orthologue *drp-1* and fusion by loss of the mitofusin orthologue *fzo-1*^15^ would impact UVB- and IR-induced stress tolerance. We found that *drp-1* was dispensable for these responses (Figure 1A and B), while lack of mitochondrial fusion (mito-fusion) strikingly sensitized the animals to stress following treatment with UVB but not IR (Figure 1A and B), suggesting the mito-fusion is essential for the maintenance of systemic homeostasis following UVB exposure. Consistently, UVB but not IR induced persistent mitochondrial fragmentation in transgenic animals expressing GFP labelled mitochondria^16^ (Figure 1C and S1B), which was exacerbated by RNAi-mediated knock down of *fzo-1* (Figure S1C). We thus found that UVB treatment leads to fragmentation of mitochondrial network, which is mitigated by the fusion machinery to maintain healthy homeostasis.

**Figure 1.**
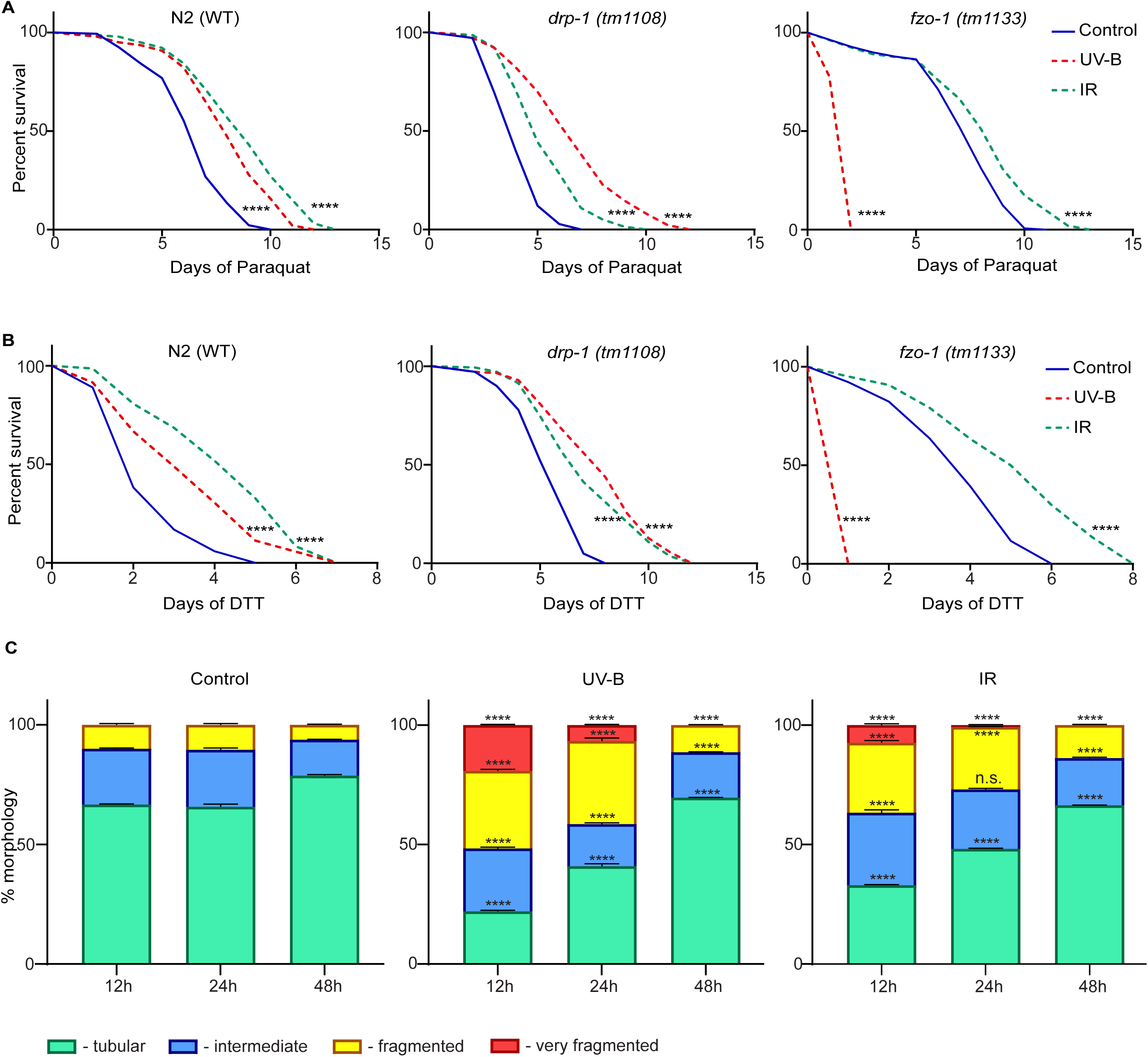
Adaptive benefits of moderate UVB exposure require mitochondrial fusion. Wild-type (N2 Bristol strain), *drp-1(tm1108),* and *fzo-1(tm1133)* nematodes were pre- treated with UV-B (850mJ/cm^2^) and IR (90Gy) at L4 stage and 48h later transferred to (**A**) 5mM Paraquat (Sigma-856177) and (**B**) 10mM DTT (Sigma-DO632) plates; survival was scored daily. Significance was measured by Mantel-Cox test, and two-tailed *p* values were computed. Each group consisted of n≥130 worms. (**C**) Transgenic animals expressing GFP-tagged mitochondria in the body wall muscle (*myo-3p::gfpmit*) were treated with UV-B (850mJ/cm^2^) and IR (90Gy) on adulthood day 1 (AD1). The % of tubular, intermediate, fragmented, and very fragmented morphologies were scored after 12h, 24h, and 48h. Significance was measured by a two-tailed unpaired t-test (with Welch’s correction). n=60 in each condition, mean and S.E.M values are presented. The asterisks compare respective morphologies in UV- and IR-treated animals versus time point matched untreated control. * *p*<0.05; ** *p*<0.01; *** *p*<0.001; **** *p*<0.0001; n.s., not significant. Representative results of at least three independent experiments are shown in all cases.

### UVB elicits a dietary restriction mimetic response that requires mitochondrial fusion

Changes of mitochondrial dynamics have previously been linked to alterations in lipid expenditure^17, 18^, and we next used whole animal Oil Red O (ORO) staining^19^ to test if mitochondrial fragmentation induced by UVB has an impact on lipid homeostasis of *C. elegans*. We found that in wild type animals UVB but not IR indeed induces a transient decline of whole body lipid content, which bares similarity to effects of dietary restriction (DR)-mimetic drugs^9^ and recovers to baseline levels within 24h post exposure (Figure 2A and 2B). Importantly, lipid levels did decline but did not recover in UVB exposed *fzo-1* mutant animals (Figure 2C and S2A), in line with the key roles of mitochondrial fragmentation and fusion in the newly uncovered DR-like effect of UVB. Importantly, at higher UVB doses the recovery of lipid levels following UV exposure was impaired also in wild type animals (Figure S2B and S2C), clearly showing that a restricted range of UV doses is effective in eliciting the DR-like effects without permanent distortion of metabolism. In line with the ability of UVB to cause the DR-like impact on the systemic metabolism, we found that UVB elicits a transient decline in whole body ATP levels in wild type animals (Figure 2D), and this effect was protracted in mitofusin mutants (Figure 2E) similar to their persistent loss of lipid storage.

**Figure 2.**
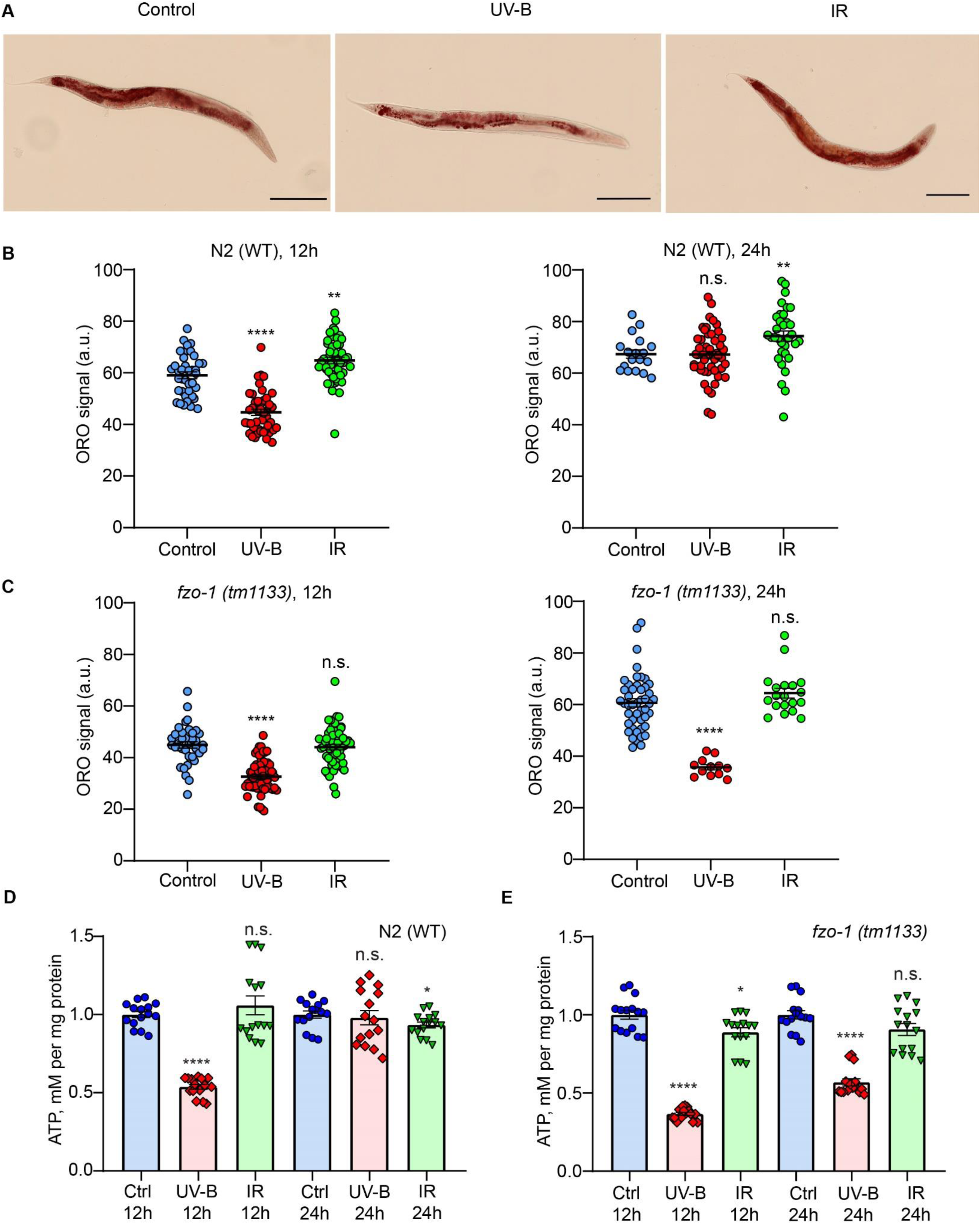
Mitochondrial fusion sustains metabolic plasticity following UVB exposure. (**A**) Representative images of Oil Red O (ORO) staining of nematodes treated with UV-B (850mJ/cm^2^) and IR (90Gy). Scale bar is 200µm. (**B**) N2 wild-type and (**C**) *fzo-1(tm1133)* strains were treated with UV-B and IR on adulthood day 1 (AD1) and ORO staining was performed after 12h and 24h. Arbitrary units (a.u.) were assigned as described in the methods section. The asterisks assess differences between treated and control groups at each time point. n≥12 for each condition and mean and S.E.M values are presented. Two-tailed unpaired t-test (with Welch’s correction) was used for statistical analysis. Whole-body ATP levels were measured in (**D**) wild-type and (**E**) *fzo-1(tm1133)* mutant strains treated with UV-B (850mJ/cm^2^) and IR (90Gy) at young (non-gravid) adult stage with samples collected at 12h and 24h post-treatment. Values are relative to respective time-matched controls. The asterisks compare treated and control groups at each time point and n=250 for each condition. Mean and S.E.M values are presented. For statistics, a two-tailed unpaired t-test (with Welch’s correction) was used. * *p*<0.05; ** *p*<0.01; *** *p*<0.001; **** *p*<0.0001; n.s., not significant. Representative results of at least three independent experiments are shown in all cases.

We next performed proteomics analysis in wild type and mitofusin mutant animals to test for molecular activities associated with dietary restriction and metabolic rewiring^9^. Consistent with the DR-like effects of UVB, we indeed found upregulation of glycolysis (Figure 3A, Table S1), downregulation of lipid droplet components (Figure 3B, Table S1) and upregulation of beta oxidation enzymes both mitochondrial (Figure 3C, Table S1) and peroxisomal (Figure 3D, Table S1) in wild type animals exposed to UVB. Strikingly, all of these rewiring activities were abrogated and even reversed in UVB-treated mitofusin mutant animals (Figure 3A-D, Table S1), in line with the key role of mitochondrial fusion in the recovery of metabolic homeostasis following UVB exposure.

**Figure 3.**
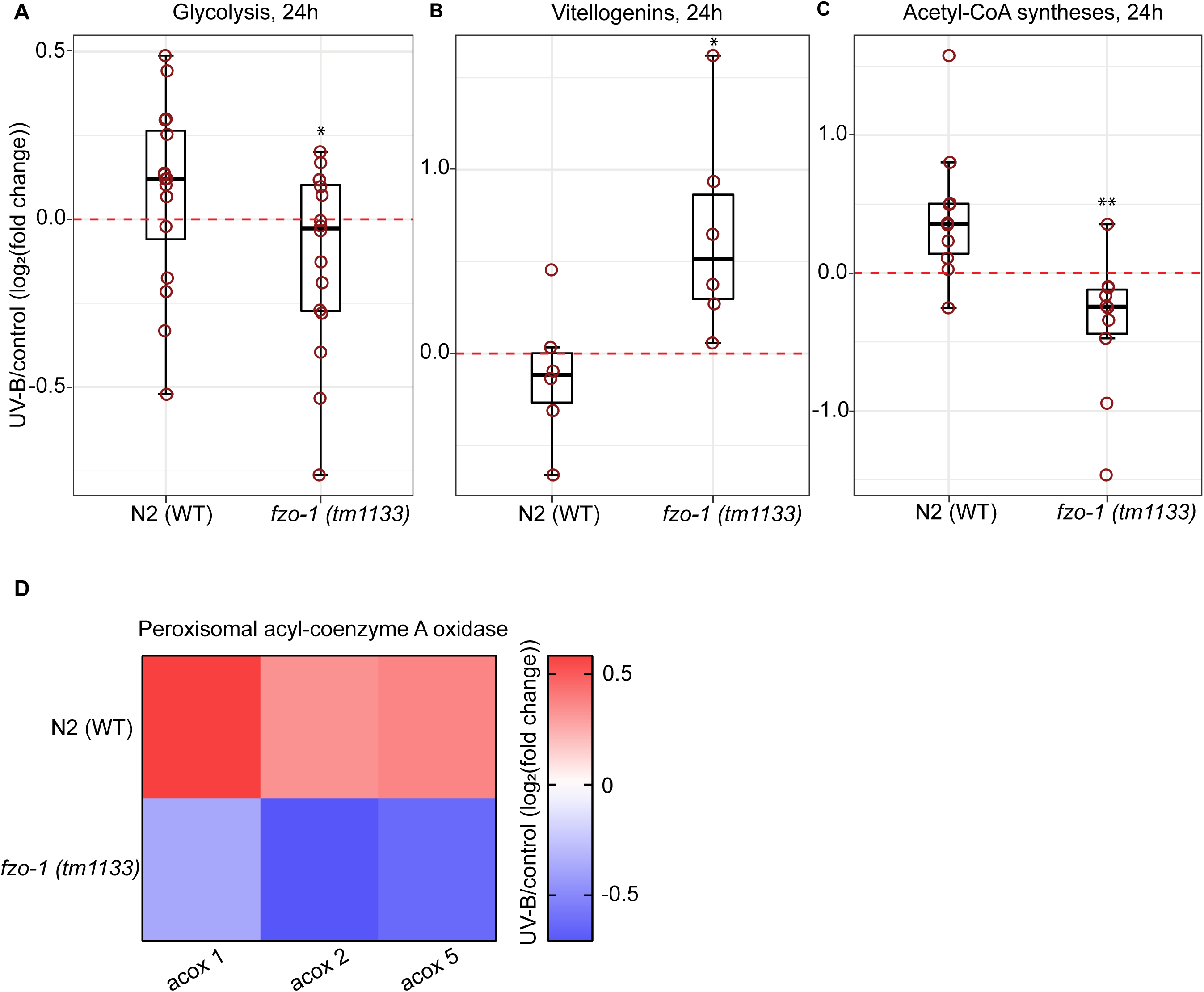
UVB treatment elicits dietary restriction mimetic effects. Wild type (N2) and *fzo-1(tm1133)* mutant nematodes were treated with 300mJ/cm^2^ UVB on adulthood day 1 and protein samples were collected after 24h. Lower UV dose was experimentally determined to avoid loss of *fzo-1* mutants due to direct UV toxicity. Box plots show average Log2 expression fold changes of (**A**) selected proteins in the glycolysis pathway, (**B**) vitellogenins and (**C**) mediators of mitochondrial beta-oxidation. In all cases, fold changes are calculated between UV-treated versus respective time point matched untreated control samples. Red circles represent individual proteins and the median fold change of the proteins belonging to each group is shown as a bold line. The first and third quartile are indicated by upper and lower limits of the boxplot and whiskers extend to 1.5 times the interquartile range from the limits of the box. Four independent pools of n=800 worms were analyzed for each condition. Wilcoxon rank-sum test and two-tailed *p* values were used for statistics. Asterisks highlight differences between respective *fzo-1(tm1133)* and wild-type N2 samples. * *p*<0.05; ** *p*<0.01; *** *p*<0.001; **** *p*<0.0001; n.s., not significant. (**D**) A heatmap of selected proteins involved in peroxisomal beta-oxidation is shown. The color bar depicts Log2 (fold change) values. Individual protein changes and respective statistics for (**A-D**) are shown in Table S1.

### Mitochondrial effects of UVB are independent of nuclear DNA damage

While both UVB and IR are inducers of nuclear DNA damage^20, 21^, the animals, which lack mitochondrial fusion, demonstrated impaired survival in response to UV but not IR (Figure 1A and B). Consistently, prominent DR-mimetic metabolic changes were seen upon UV but not IR exposure (Figure 2A-E). Collectively these findings indicate that the DR-like effect of UV is likely independent of its ability to inflict nuclear genotoxic stress. In previous work, we showed that nuclear DNA damage triggered by UVB and IR was linked to systemic stress tolerance by elevated innate immune signaling and enhanced proteostasis^7^. While both responses were detected in our current proteomics profiles of UVB-exposed *C. elegans*, they were not abrogated in mitofusin mutant animals (Figure S3A and B, Table S2) again suggesting independence of nuclear and mitochondrial effects of UVB.

The DNA helix distorting lesions induced by UVB are repaired by Nucleotide Excision Repair (NER) pathway, and such lesions accumulate in NER mutant animals^21^. Interestingly, previous study showed that congenital defects in Transcription Coupled NER led to mitochondrial hyper-fusion rather than fragmentation in the absence of UVB exposure^22^. To complement this published finding, we inactivated the second NER branch – Global Genome (GG) NER by treating L1 larva with RNAi against the GG NER component *rad-23*^21^, followed by UVB exposure and assessment of mitochondrial morphology. We found elevated baseline numbers of fragmented mitochondria in control GG NER-deficient worms (Figure S3C) in line with the recently reported strong effects of persistent DNA damage on metabolic homeostasis^12^. However, an increase of mito- fragmentation upon UVB exposure and its timely recovery within 48h could still be observed in *rad-23* knock down animals (Figure S3C), suggesting, together with the above evidence, that mitochondrial stress induced by UVB is independent of its ability to trigger nuclear DNA damage.

### DR-mimetic responses to UVB are not linked to lasting mitochondrial damage in wild type animals

We next asked if UVB disturbs mitochondrial homeostasis by inflicting lasting damage of mitochondrial DNA and/or mitochondrial proteins. Both these damages converge in triggering mitochondrial unfolded protein response (UPR MT), which can be visualized *in vivo* in *hsp-6p::GFP* reporter nematodes^23^. We found however that neither UVB nor IR triggered detectable UPR MT activity at their beneficial doses used in this study (Figure 4A and S4). Consistently, proteomics analysis showed no reduction of mitochondrial content in UVB-exposed wild type animals at either 24h or 12h post treatment (Figure 4B, Table S3 and S5A, Table S5), and mutants lacking key mitophagy mediator PINK-1^24^ did not fail in developing UVB- and IR-induced stress tolerance (Figure 4C). Collectively, these results reveal that moderate UVB treatment does not cause lasting mitochondrial damage and does not trigger significant mitochondrial turnover by mitophagy in wild type animals. Conversely, in mitofusin mutants UVB treatment led to significant and persistent decline of mitochondrial content (Figure 4B, Table S3 and S5A, Table S5), which was accompanied by strong upregulation of the V-ATPase subunits required for lysosomal acidification during autophagy and mitophagy^25, 26^ (Figure 4D,Table S3 and S5B, Table S5), altogether demonstrating that mitochondrial fusion is protective against lasting negative impact of UVB on mitochondrial integrity.

**Figure 4.**
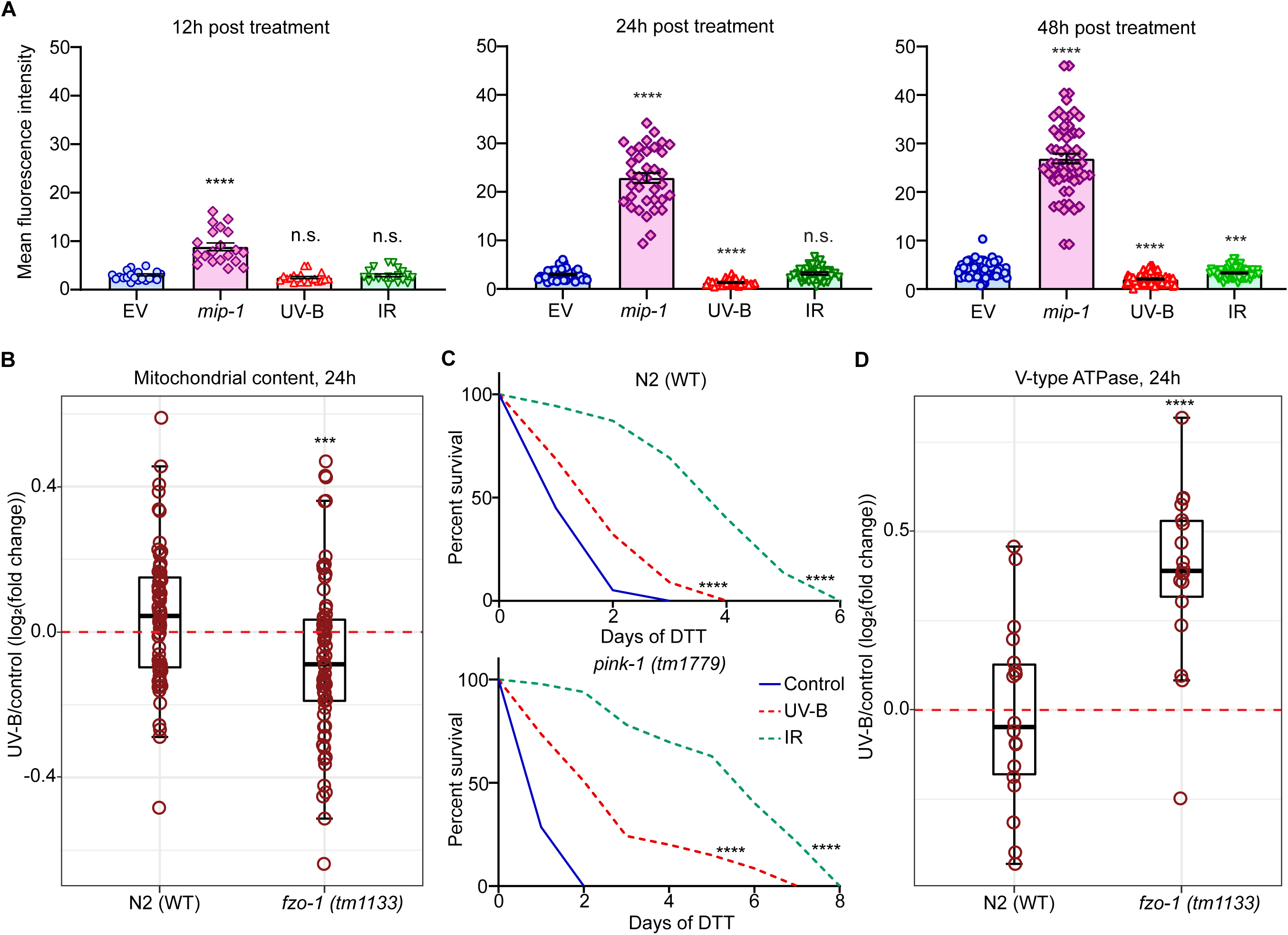
Moderate UVB exposure does not trigger lasting mitochondrial damage in wild type animals. (**A**) Transgenic animals expressing GFP under the promoter of the *hsp-6* mitochondrial chaperone gene were treated with UV-B (850mJ/cm^2^) and IR (90Gy) at young (non- gravid) adult stage, and GFP fluorescence was quantified after 12h, 24h, and 48h. *mip- 1* RNAi was used as a positive control for UPR^MT^ induction. n≥20 worms were analyzed for each condition. Significance was measured by a two-tailed unpaired t-test (with Welch’s correction); mean and S.E.M values are presented. Asterisks compare treated groups to time point matched untreated controls. Box plots show average Log2 expression fold changes of (**B**) selected mitochondrial ribosome proteins and (**D**) V-type ATPase components. Fold changes were calculated between UV-B treated samples and timepoint matched untreated controls. Four independent pools of n=800 worms were collected at 24h post-exposure to 300mJ/cm^2^ UV-B on adulthood day 1 (AD1). Red circles represent individual proteins, and box plot parameters are as described in Figure 3 (A- C). Wilcoxon rank-sum test and two-tailed *p* values were used for statistical analysis. Asterisks compare *fzo-1(tm1133)* values to corresponding wild-type N2 controls. (**C**) Wild- type, and *pink-1(tm1779)* strains were pre-treated with UV-B and IR at L4 stage and after 48h transferred to 10mM DTT (Sigma-DO632) plates, and survival was scored daily. Significance was measured by Mantel-Cox test, and two-tailed *p* values were computed. Each group consisted of n≥130 worms. * *p*<0.05; ** *p*<0.01; *** *p*<0.001; **** *p*<0.0001; n.s., not significant. Representative results of at least three independent experiments are shown in **A** and **C**. Individual protein changes and respective statistics for **B** and **D** are shown in Table S3.

### Homeostatic recovery upon UVB exposure requires mitochondrial biogenesis and Ca^2+^ signaling

By dedicated analysis of the mitochondrial proteome following UVB exposure, we found that the expression of electron transport chain (ETC) components^9^ was surprisingly elevated in UV treated mitofusin mutants (Figure 5A, Figure S5C, Tables S4-S5) despite the overall decline of mitochondrial content seen in these animals (Figure 4B, Table S3 and S5A, Table S5). This phenotype held true for each individual ETC complex with highest significance seen for complexes I and V, possibly due to bigger number of components detected for these complexes (Figure S6A-E, Table S6). This observation suggested that UVB-induced mitochondrial distortions might trigger the biogenesis of ETC-enriched organelles that are integrated into the network via fusion to restore metabolic equilibrium and mitochondrial integrity. In *C. elegans,* mitochondrial biogenesis is controlled by the conserved transcription factor SKN-1/Nrf2^24^, and inactivation of this gene by mutation or RNAi indeed prevented mitochondrial network recovery following UVB exposure (Figure S7A) and sensitized the worms to UVB toxicity (Figure 5B and S7B). Because SKN-1 is known to respond to an increase of cytosolic Ca^2+^ levels triggered by mitochondrial dysfunction^24, 27^, we next exposed the animals to Ca^2+^ chelator EGTA^24^ and indeed found that Ca^2+^ removal prevented the recovery of mitochondrial integrity in UVB-exposed animals similar to phenotypes seen upon RNAi-mediated inactivation of *skn-1* and *fzo-1* (Figure S7C). Consistently, our tests in transgenic animals expressing Ca^2+^ sensor GCaMP3^28^ detected *in vivo* Ca^2+^ release at 12h post UVB exposure (Figure 5C). Moreover, UVB-treatment of human primary skin fibroblasts (a relevant cell type with regard to UV effects in humans^29^) did not lead to lasting disruption of mitochondrial integrity unless it was accompanied by EGTA exposure, in which case UV promoted loss of mitochondrial membrane potential (MMP) and cell death (Figure 5D and S8).

**Figure 5.**
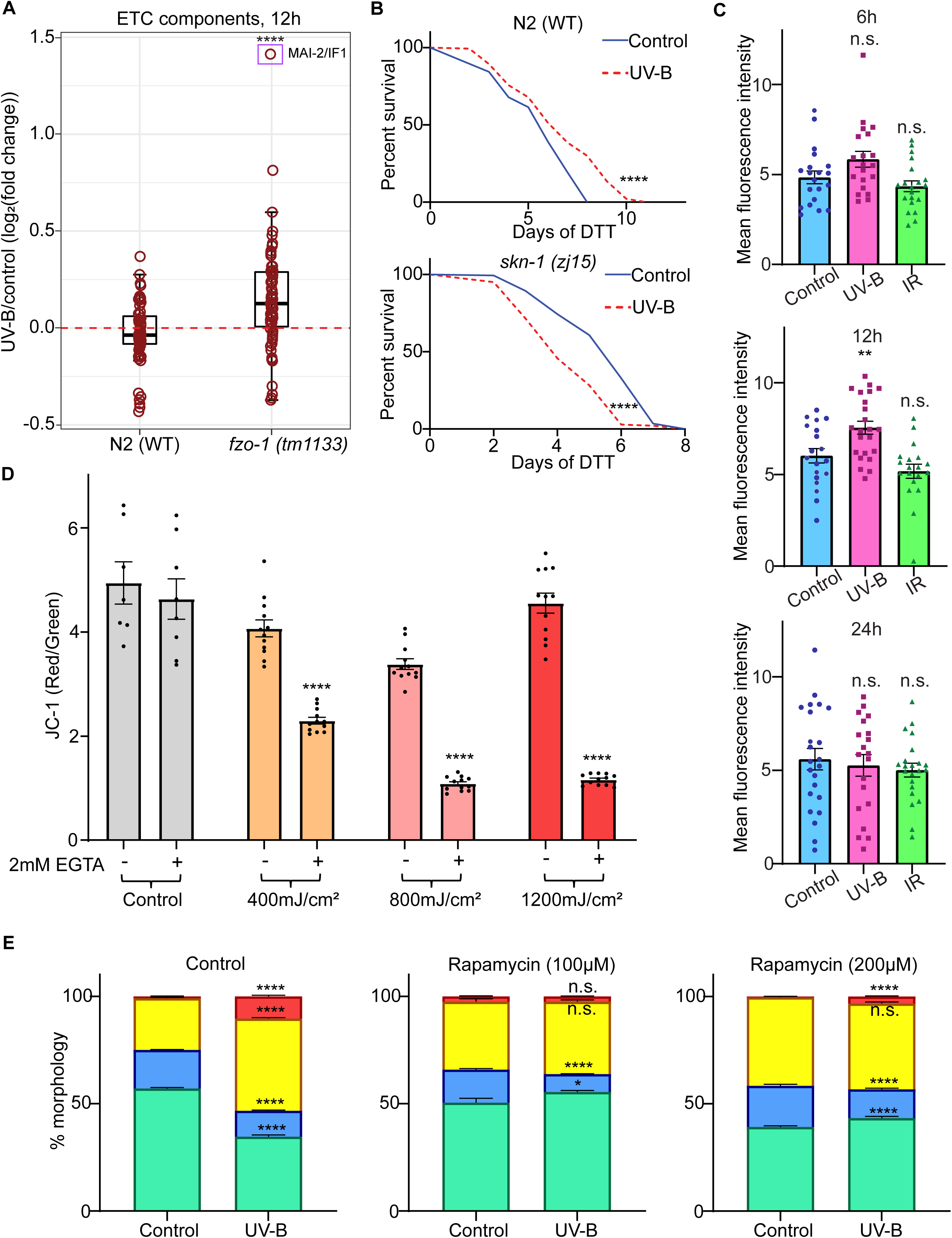
Adaptive UVB effects involve ΔΨ distortion, mitochondrial biogenesis and Ca^2+^ signaling. Protein samples were collected as described in Figures 3 and 4. (**A**) Box plots show average Log2 expression fold changes of selected ETC components at 12h post exposure to 300mJ/cm^2^ UV-B. Fold changes were calculated in UV-B treated versus time point matched control groups. Four independent pools of n=800 worms were collected and analyzed for each condition. Red circles represent individual proteins, and box plot parameters are as described in Figure 3 (A-C). Wilcoxon rank-sum test and two-tailed *p* values were used for statistics. Asterisks compare *fzo-1(tm1133)* and corresponding wild- type N2 samples. Purple rectangle highlights MAI-2/IF1 protein. (**B**) Wild-type (N2 Bristol strain) and *skn-1(zj15)* mutant animals were pre-treated with UV-B (850mJ/cm^2^) at L4 stage and after 48h transferred to 10mM DTT (Sigma-DO632) plates; survival was scored daily. Significance was measured by Mantel-Cox test, and two-tailed *p* values were computed. Each group consisted of n=140 worms. (**C**) Transgenic animals expressing calcium sensor GCaMP3 in body wall muscle were treated with UV-B (850mJ/cm^2^) and IR (90Gy) at young (non-gravid) adult stage, and fluorescence was quantified after 6h, 12h, and 24h. n≥20 worms were analyzed for each condition. Significance was measured by a two-tailed unpaired t-test (with Welch’s correction); mean and S.E.M values are presented. Asterisks compare treated and untreated groups at each time point. (**D**) PD41 human skin fibroblasts were pre-treated with 400 mJ/cm^2^, 800 mJ/cm^2^ and 1200 mJ/cm^2^ of UV-B, and later incubated in presence of 2mM EGTA. Mitochondrial membrane potential was measured by JC-1 assay after 24h. Significance was assessed by a two- tailed unpaired t-test (with Welch’s correction); mean and S.E.M values are presented. Asterisks compare respective EGTA plus and minus conditions. (**E**) (*myo-3p::gfpmit*) transgenic animals expressing GFP-tagged mitochondria in the body wall muscle were pretreated with indicated amounts of Rapamycin (R-5000, LC laboratories) for 24hours before UV-B (850mJ/cm^2^) treatment on adulthood day 1 (AD1). The % of tubular, intermediate, fragmented, and very fragmented mitochondria were scored after 12h. Significance was measured by a two-tailed unpaired t-test (with Welch’s correction). n=60 in each condition, mean and S.E.M values are presented. Asterisks compare respective UV-B treated and untreated groups. * *p*<0.05; ** *p*<0.01; *** *p*<0.001; **** *p*<0.0001; n.s., not significant. In **B-D** representative results of at least three independent experiments are shown. Individual protein changes and respective statistics for **A** are shown in Table S4.

We next reasoned that coordinated upregulation of ETC components in UV treated animals (Figure S6A-E, Table S6) could represent a compensatory response to the direct distortion of oxidative phosphorylation (OXPHOS) by UVB. Interestingly, early studies in isolated mitochondria indeed suggested that UV is able to impair OXPHOS^30, 31^ and disrupt mitochondrial proton gradients, which in turn can trigger mito-fragmentation^32^ and Ca^2+^ release^33^. This negative impact of UV on the mitochondrial bioenergetics could be mitigated by provision of ATP to the organelles^34^, consistent with the capacity of complex V to act as a proton pump for ΔΨ stabilization by switching from ATP synthesis to ATP hydrolysis^35, 36^. Strikingly, the highest upregulated respiratome component in UVB- exposed mitofusin mutants was MAI-2 (Figure 5A,Table S4 and S5C,Table S5), the *C. elegans* ortholog of the inhibitory factor 1 (IF1)^37^, and MAI-2 upregulation upon UV was seen only in the mutants and not in WT animals (Figure S5D). The role of IF1 is to inhibit the ATP hydrolysis function of complex V preventing mitochondria from causing whole cell ATP exhaustion under conditions of persistent OXPHOS failure^38^. Consistently, stabilization of the *in vivo* ATP content by rapamycin exposure^9^ alleviated UV-induced mitochondrial fragmentation, in line with OXPHOS distortion being the primary trigger of this phenotype (Figure 5E).

Collectively, our data supports the model that distortion of mitochondrial OXPHOS by moderate UVB exposure triggers mitochondrial fragmentation and elicits a Ca^2+^ signal that initiates biogenesis of ETC-enriched mitochondria followed by their integration into the network via fusion. This recovery process prevents lasting mitochondrial damage in UV exposed cells, while the initial decline of mitochondrial output triggered by UVB elicits DR-like metabolic rewiring and contributes to systemic stress tolerance (Figure S9).

### Aging-linked decline of mitochondrial fusion sensitizes old animals to acute UVB toxicity

So far, we found that moderate doses of UVB trigger DR-mimetic metabolic changes by inducing mitochondrial fragmentation, and mito-fusion is required to ensure the transient nature of these effects avoiding lasting mitochondrial damage and UVB toxicity. Concurrently, previous studies demonstrated a decline of mito-fusion during aging^9, 39^. We thus asked if aging interferes with the recovery of the animals from metabolic stress induced by UVB. Indeed, while mitochondrial fragmentation was transient in UVB- exposed young nematodes, it was more pronounced and more persistent in the old ones (Figure 6A). Consistently, while young animals benefited from UVB exposure by developing adaptive stress tolerance (Figure 6B), the survival of old UVB-treated nematodes was significantly compromised demonstrating UVB toxicity (Figure 6C). Our data thus indicates that aging-associated decline of mitochondrial fusion disables the metabolic benefits of UVB in old animals sensitizing old organisms to UVB toxicity in a manner, which resembles the defective responses of mitofusin mutants to UVB (Figure 6D).

**Figure 6.**
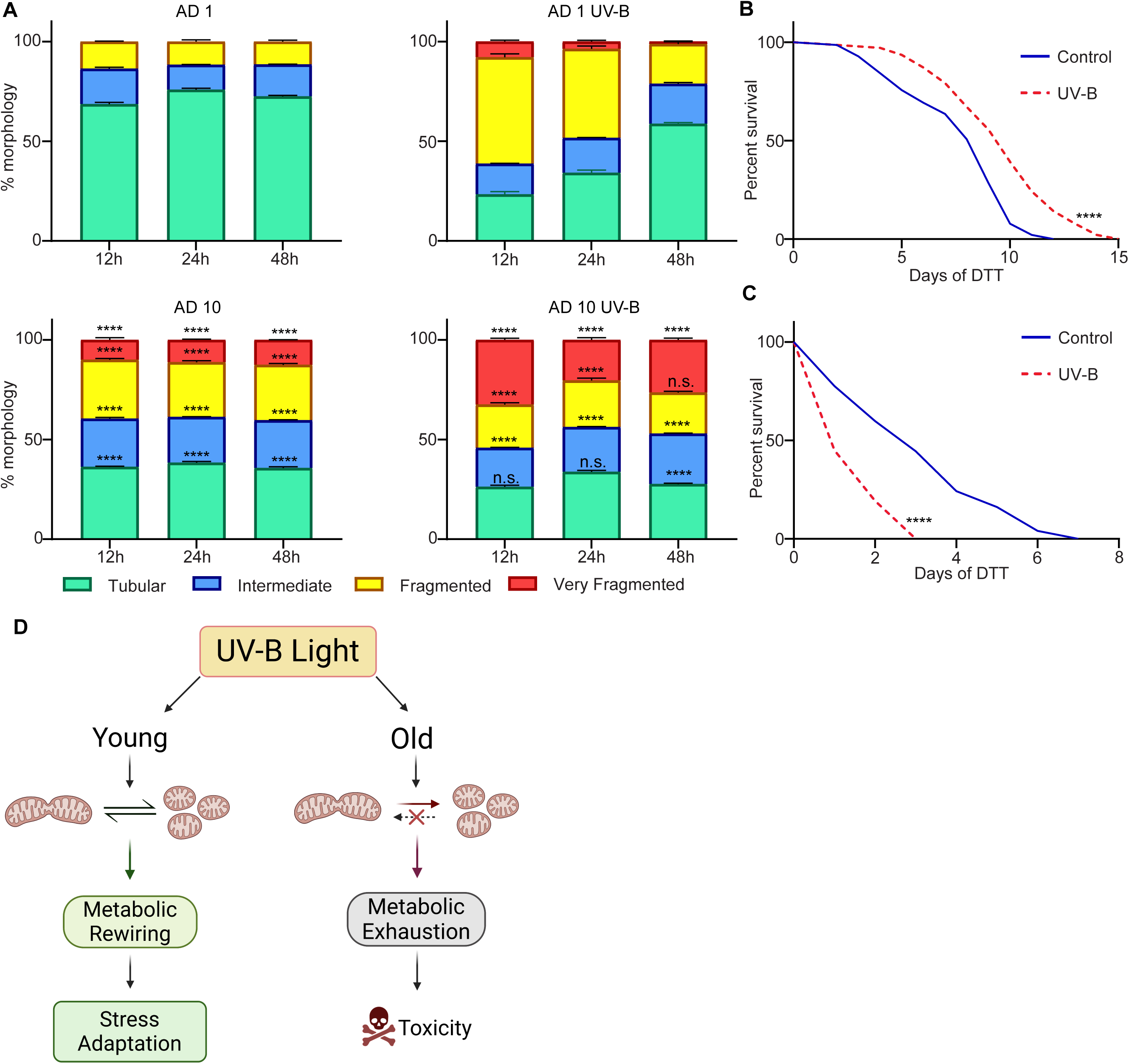
Aging-associated impairment of mito-fusion abrogates UVB benefits and triggers UVB toxicity. (**A**) Young (adulthood day 1, AD1) and old (AD10) transgenic animals expressing GFP tagged mitochondria in the body wall muscle (*myo-3p::gfpmit*) were treated with UV-B (850mJ/cm^2^). The % of tubular, intermediate, fragmented, and very fragmented mitochondria were scored after 12h, 24h, and 48h. Significance was measured by a two- tailed unpaired t-test (with Welch’s correction), n=60 in each condition, mean and S.E.M values are presented. The asterisks compare respective morphologies of old nematodes with time point- and treatment-matched young control. Young (AD1) (**B**) and old (AD10) (**C**) wild-type animals were treated with UV-B and after 48h transferred to DTT (Sigma-DO632) plates; survival was scored daily. Significance was measured by Mantel- Cox test, and two-tailed *p* values were computed. At least three independent experiments were conducted in each case, (**B**) n=140 and (**C**) n≥60 for each experimental condition. * *p*<0.05; ** *p*<0.01; *** *p*<0.001; **** *p*<0.0001; n.s., not significant. (**D**) Model: in young animals, efficient mitochondrial fusion supports metabolic recovery after UV exposure, promoting homeostatic benefits, while aging-associated fusion defects sensitize old organisms to UV toxicity. The image is created with BioRender (biorender.com).

### UVB irradiation of the skin elicits body-wide DR-mimetic phenomena in young mice

While penetration of the whole body with UVB is feasible in small and optically transparent *C. elegans*, in mammals UVB light cannot penetrate beyond the superficial skin layers^40^. We next performed proof-of-concept experiments in young mice to test if UVB irradiation of the skin is sufficient for triggering systemic responses that resemble fasting and/or DR. Interestingly, we found that irradiation of the approx. 1/3 of the mouse skin surface (in the back area and following hair removal) with a single UVB dose of 100 mJ/cm^2^ caused fasting-like changes in distant mouse tissues (Figure S10A-C). For instance, transient cell size shrinkage was seen in the abdominal white adipose tissue within hours of UVB treatment consistent with the fatty acid release from these cells as in fasting^41^ (Figure S10B). At the same time, a transient increase of lipid levels was detected in the liver of UVB-exposed mice similar to changes that accompany the process of ketogenesis during fasting^42, 43^. While additional molecular assays are needed to comprehensively liken the observed UVB-induced phenomena to DR and fasting, our first observations suggest that UVB irradiation may be applicable as a non-pharmacological DR mimetic treatment in mammals. Of note the dose we used on mouse skin was below the range of UVB doses (200-800 mJ/cm^2^) routinely given in humans for the treatment of skin diseases such as atopic dermatitis and psoriasis^44^, and we didn’t observe obvious signs of radiation damage (e.g. erythema or sunburn) in the UV-treated skin areas of mice during the entire observation period of 36h (data not shown).

## Discussion

While positive effects of moderate UV exposure on organismal health are commonly known, the cellular and systemic basis of these benefits remains elusive. In this study, we used *C. elegans* and human skin fibroblasts to illuminate the ability of UVB to alter mitochondrial bioenergetics with significant cellular and systemic consequences. We show that UVB irradiation triggers a decline of mitochondrial membrane potential leading to a rapid fragmentation of the mitochondrial network, reduction of the systemic ATP output, and swift metabolic rewiring that closely resembles effects of dietary restriction at the molecular and organismal levels. The UVB-induced disruption of mitochondrial integrity is followed by a recovery process mediated by Ca^2+^, SKN-1/Nrf2 and mitochondrial fusion to protect cells from UV toxicity and restore the metabolic balance (Figure S9). Our findings thus lay out a new mechanistic connection between UVB and cellular and systemic metabolism that is executed via changes of mitochondrial respiration and network integrity. Importantly, UVB-induced mitochondrial fission and distorted respiration observed in our study are opposite of mitochondrial effects (network protection and OXPHOS enhancement) of the vitamin D^45–47^. Vitamin D synthesis is triggered by the UV irradiation of the skin with known impact on the systemic homeostasis and metabolism^47^. However, several metabolic UV effects such as prevention of obesity in high fat diet exposed mice, could be achieved also in animals deficient in vitamin D signaling^4, 6^. In this context, our study reveals for the first time a potential mechanism of the vitamin D-independent effects of UV light on the systemic fat storage. Interestingly, anti-obesity effects of the skin sunlight exposure (including a natural UVB component) were observed also in humans^48^, and the UV-triggered mechanism we found is clearly conserved between nematodes and human skin cells. Because skin is a large organ, and its responses are sufficient to generate systemic organismal changes^3, 5^, it is feasible that the newly discovered DR-mimetic effect of UVB can even be translationally explored, especially given the compelling evidence from the mouse models^4, 6^. Moreover, UVB exposure of limited skin areas is already applied in humans as a safe and effective therapy against psoriasis and atopic dermatitis (eczema)^49, 50^. We therefore conducted a pre-clinical proof-of-concept study in young mice and intriguingly observed that the irradiation of the skin with a single low-to-moderate dose of UVB elicits a whole organism fasting-like response in the ad-libitum fed animals. Finally, we identified aging-associated decline of mitochondrial fusion as a cause of metabolic and cellular UVB toxicity that likely poses risks for older individuals given the omnipresence of sunlight in the daily life of humans. Collectively, our study describes novel cellular targets of UV irradiation and illuminates new mechanistic links between an environmental stimulus (UVB) and systemic metabolism. Our findings suggest that UVB exposure of the skin may hold potential as a fast and effective non-pharmacological DR mimetic treatment, while mitochondrial aging is revealed as a likely confounding factor of the adverse UV effects in the elderly.

## Supporting information

Statistics source data

## Acknowledgements

We thank the Proteomics and Imaging Core Facilities, and Gamma Irradiation Core Service at FLI for supporting this study. The FLI is a member of the Leibniz Association and is financially supported by the Federal Government of Germany and the State of Thuringia. ME is supported by the ERC CoG LifeLongFit of the European Commission, the Carl-Zeiss-Stiftung via IMPULS consortium and is a member of Cluster of Excellence Balance of the Microverse of the Jena University funded by the Deutsche Forschungsgemeinschaft (DFG). AM received support from the Professor Doktor Dieter Platt Stiftung (Research award 2022) and the German Research Council (Deutsche Forschungsgemeinschaft, DFG) via the Research Training Group Adaptive Stress Responses (GRK 1715). YL is a member of the Jena-Shenzhen joint PhD program and receives the respective stipend.

## Author contribution

ME conceptualized and designed the study; AM, YW and SL performed experiments in nematodes and human cells; ME, AM analyzed the data; AM prepared figures and performed statistical analysis; YL performed proof-of-concept mouse experiments and LF supervised the mouse tests; ME wrote the manuscript and AM, YL and LF reviewed the manuscript.

## Declaration of interests

ME and AM are listed as inventors on a patent application based on some of the work described here.

## Data availability

The complete proteomics data set is deposited at ProteomeXchange Consortium via the PRIDE partner repository with an identifier: PXD034910.

---

**Figure S1.**
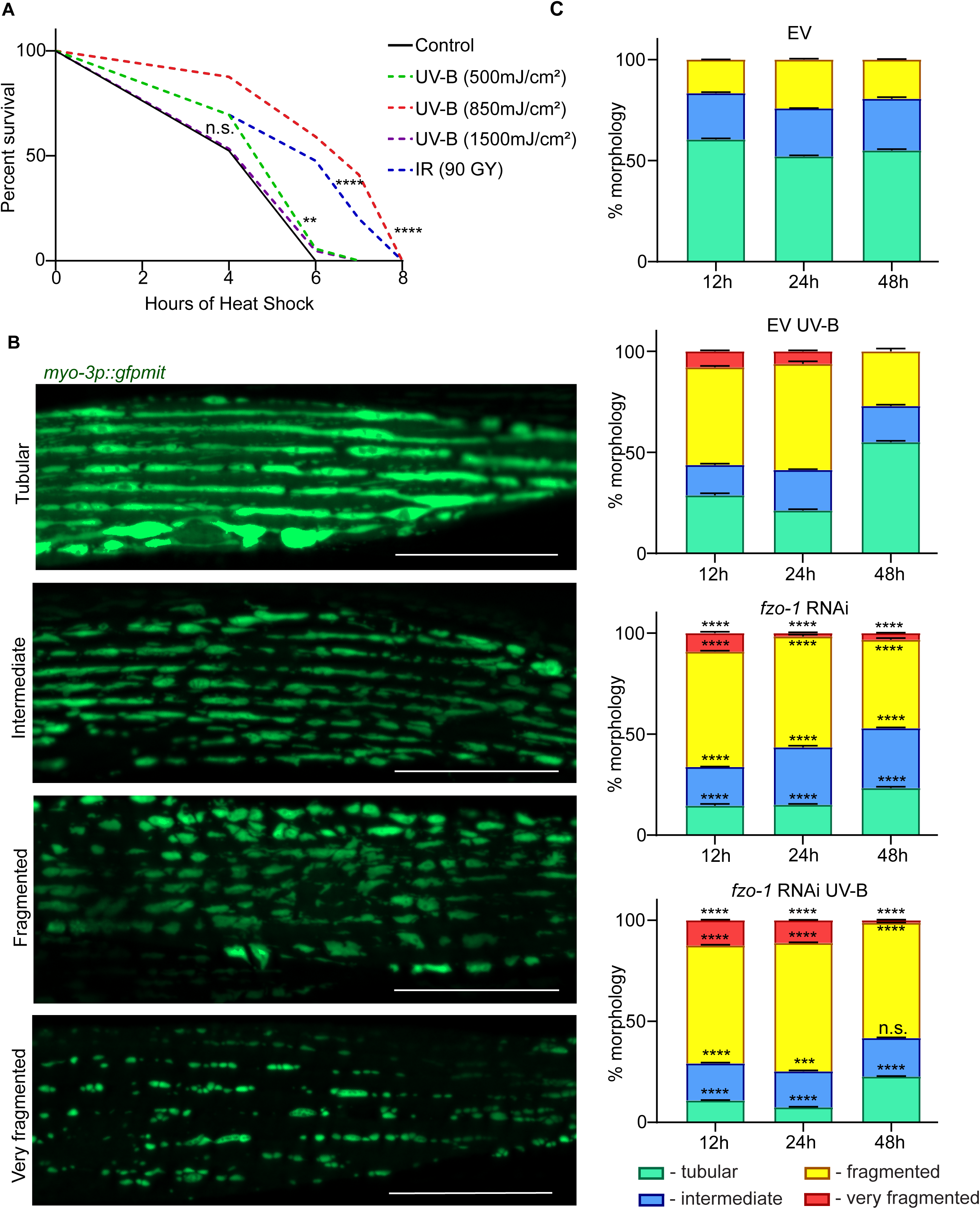
UVB-induced mitochondrial fragmentation is exacerbated by mitofusin gene knock down. (**A**) L4 stage N2 wild-type worms were treated with different doses of UV-B (500mJ/cm^2^, 850mJ/cm^2^ and 1500mJ/cm^2^) and IR (90Gy). After 48h at 20°C worms were treated with heat stress (35°C) and survival was scored at 2h, 4h, 6h, 7h, 8h post-exposure. UV-B 850mJ/cm2 and IR 90Gy were determined as optimal doses for further stress assays. Significance was measured by the Mantel-Cox test, and two-tailed *p* values were computed. At least three independent experiments were conducted in each case, n≥100 for each experimental condition. (**B**) Representative images of tubular, intermediate, fragmented, and very fragmented mitochondria in transgenic animals expressing GFP labelled mitochondria in the body wall muscle (*myo-3p::gfpmit*). The scale bar is 20µm. (**C**) (*myo-3p::gfpmit*) transgenic animals were grown on EV and *fzo-1* RNAi bacteria from the L1 stage. The nematodes were treated with UV-B (850mJ/cm^2^) on adulthood day 1 (AD1) and % of tubular, intermediate, fragmented, and very fragmented mitochondria were scored 12h, 24h and 48h post-exposure. Significance was measured by a two-tailed unpaired t-test (with Welch’s correction), n=60 in each condition, mean and S.E.M values are presented. The asterisks compare respective morphologies of *fzo-1* RNAi nematodes with time point- and treatment-matched EV control. Representative results of at least three independent experiments are shown. * *p*<0.05; ** *p*<0.01; *** *p*<0.001; **** *p*<0.0001; n.s., not significant.

**Figure S2.**
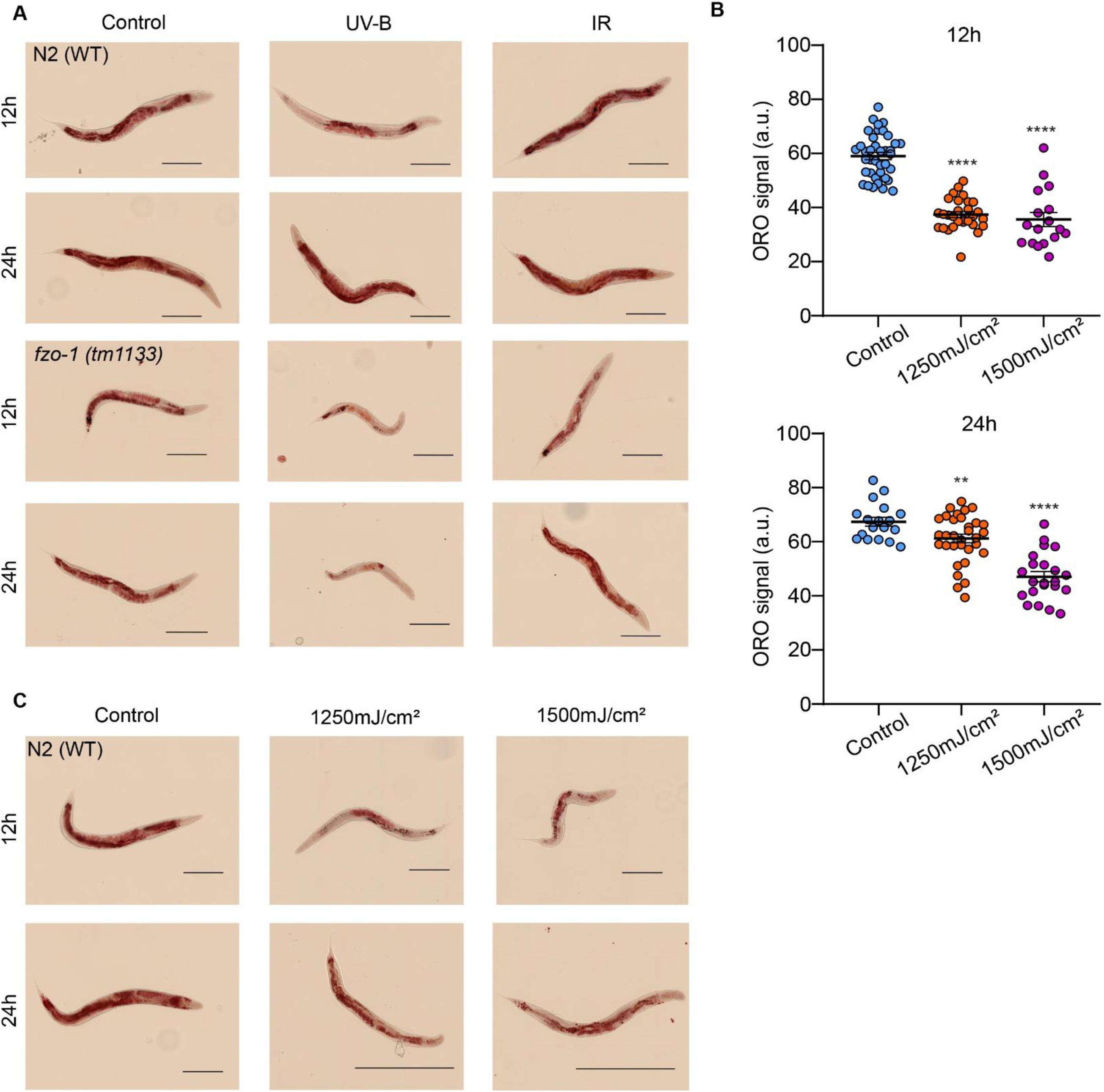
Metabolic benefits occur within limited range of moderate UVB doses. Representative images of whole-animal Oil Red O (ORO) staining of (**A**) wild-type N2 and *fzo-1(tm1133)* mutant animals at 12h and 24h after treatment with 850mJ/cm^2^ of UV-B and 90Gy IR, and (**C**) wild-type worms at 12h and 24h after treatment with 1250mJ/cm^2^ and 1500mJ/cm^2^ of UV-B; the scale bar is 200µm. (**B**) Quantification of ORO staining in N2 wild-type animals treated with 1250mJ/cm^2^ and 1500mJ/cm^2^ of UV-B at 12h and 24h post-exposure. Arbitrary units (a.u.) are assigned as described in the methods section. UV exposures for A-C were performed on adulthood day 1. Asterisks compare UV-treated and control groups at respective time points. n≥17 for each condition and mean and S.E.M values are presented. A two-tailed unpaired t-test (with Welch’s correction) was used for statistical analysis. Representative results of at least three independent experiments are shown. * *p*<0.05; ** *p*<0.01; *** *p*<0.001; **** *p*<0.0001; n.s., not significant.

**Figure S3.**
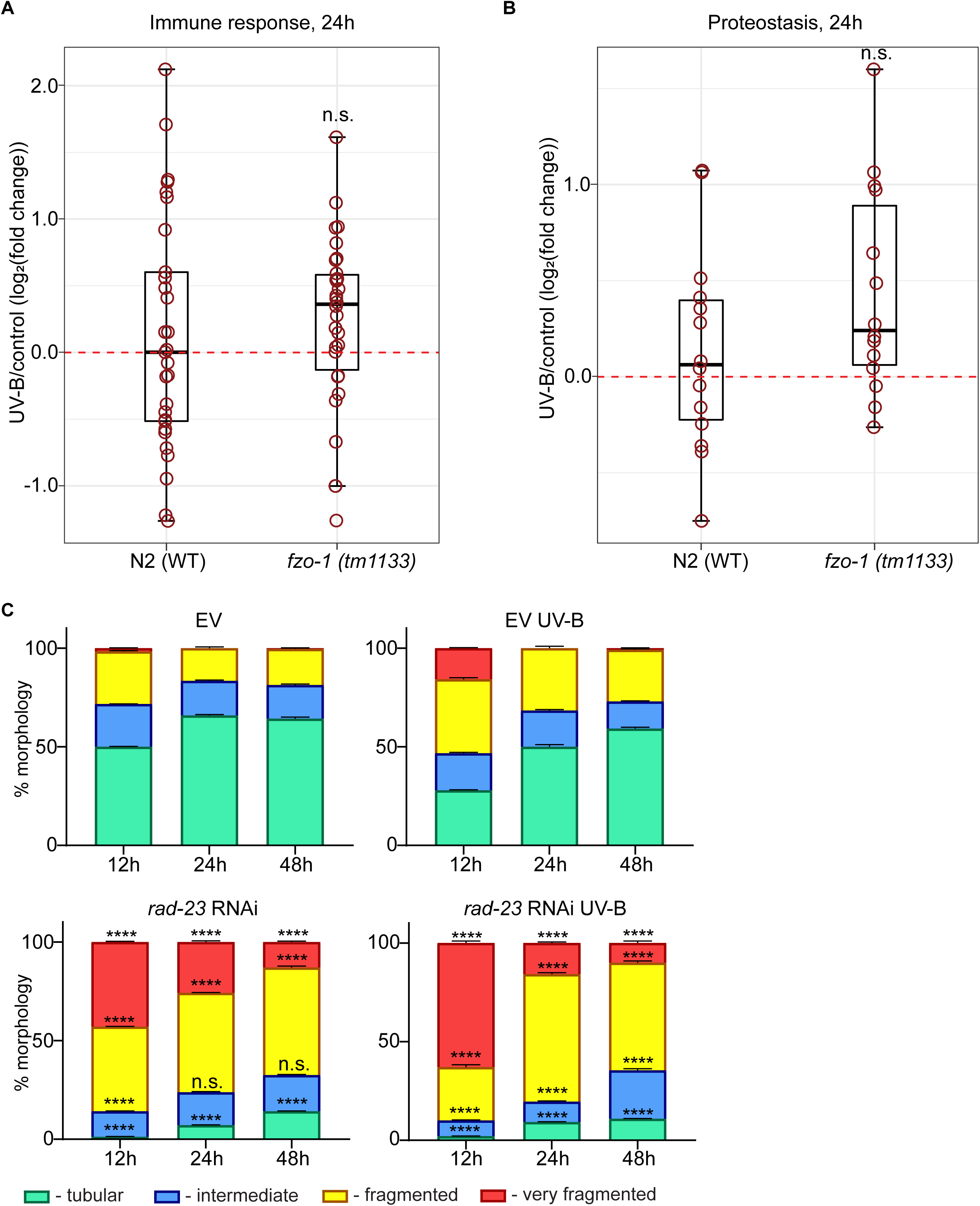
Mitochondrial and nuclear effects of UVB are independent. UV treatment and proteomics sample collection was carried out as described in Figure 3. Box plots showing average Log2 expression fold changes of selected C-type lectins (**A**) and heat shock proteins (**B**) are presented. Fold changes were calculated between UV-B treated and respective time-point matched control groups. Red circles represent individual proteins and box plot parameters are as described in Figure 3 (A-C). Four independent pools of n=800 worms were analyzed for each condition. Wilcoxon rank-sum test and two-tailed *p* values were used for statistical analysis. Asterisks compare respective *fzo-1(tm1133)* and wild-type N2 samples. (**C**) Transgenic animals expressing GFP-tagged mitochondria in the body wall muscle (*myo-3p::gfpmit*) were grown on EV and *rad-23* RNAi bacteria from the L1 stage and treated with UV-B (850mJ/cm^2^) on adulthood day 1; the % of tubular, intermediate, fragmented, and very fragmented mitochondria were scored at 12h, 24h and 48h post-exposure. Significance was measured by a two-tailed unpaired t-test (with Welch’s correction), n=60 in each condition, mean and S.E.M values are presented. The asterisks compare respective morphologies of *rad-23* RNAi nematodes with time point- and treatment-matched EV control. Representative results of at least three independent experiments are shown. * *p*<0.05; ** *p*<0.01; *** *p*<0.001; **** *p*<0.0001; n.s., not significant. Individual protein changes and respective statistics for **A** and **B** are shown in Table S2.

**Figure S4.**
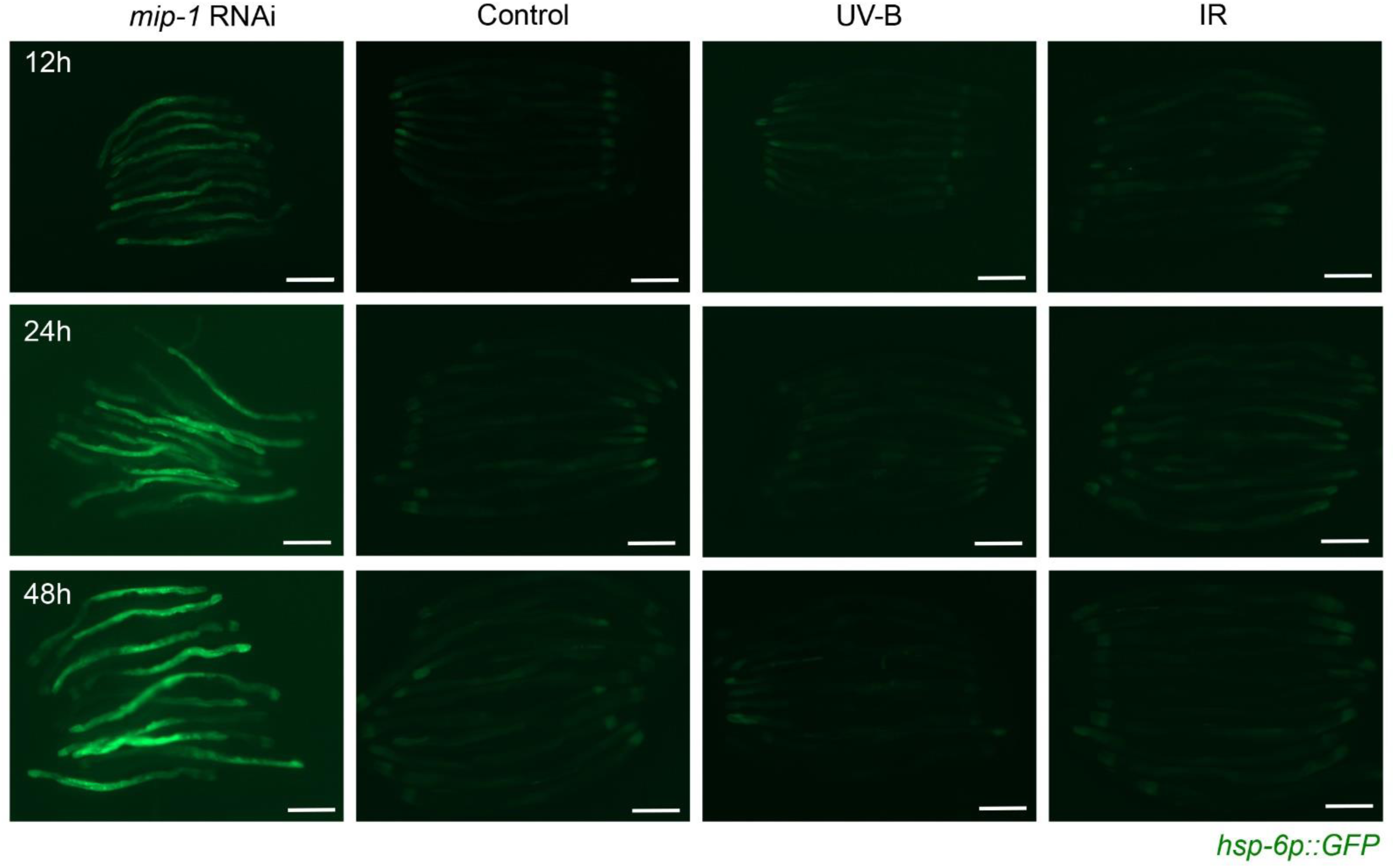
Moderate UVB and IR treatments do not activate mitochondrial UPR. Representative images of the transgenic animals expressing GFP under control of the *hsp-6* mitochondrial chaperone gene promoter and treated with UV-B (850mJ/cm^2^) and IR (90Gy) at young (non-gravid) adult stage. *mip-1* RNAi was used as a positive control that induces UPR^MT^. The microscopy pictures were taken at 12h, 24h and 48h post treatment. The scale bar is 200µm.

**Figure S5.**
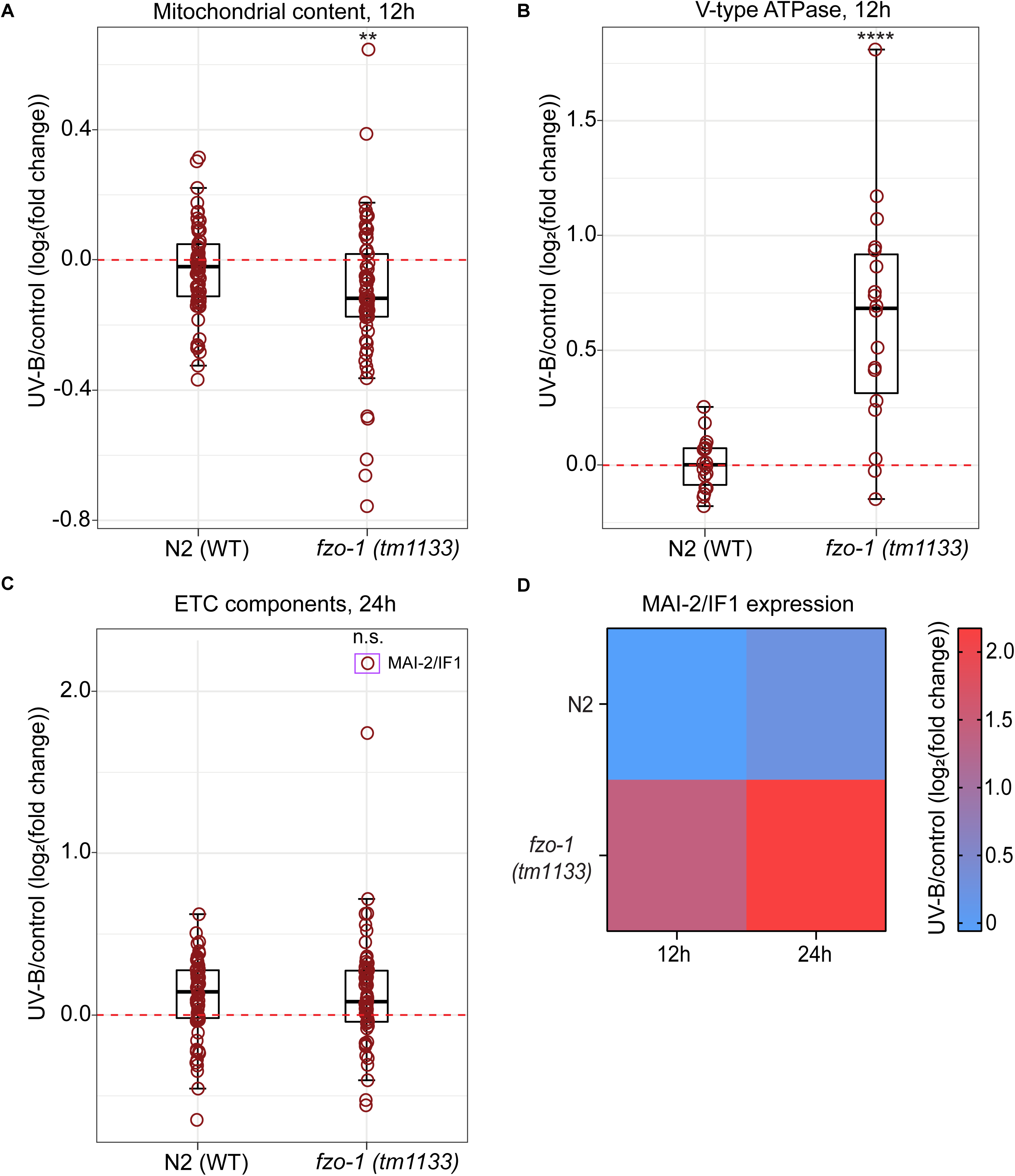
UVB exposure elicits changes of mitochondrial and lysosomal proteomes. UV treatment and proteomics sample collection was carried out as described in Figure 3. Box plots showing average Log2 expression fold changes of (**A**) selected mitochondrial ribosome proteins, (**B**) V-type ATPase components and (**C**) ETC components at indicated times following UV exposure are presented. In all cases, fold changes were calculated between UV-treated and respective time point matched control samples. In **C**, purple rectangle highlights MAI-2/IF1 protein. Red circles represent individual proteins and box plot parameters are as described in Figure 3 (A-C). Four independent pools of n=800 worms were analyzed for each condition. Wilcoxon rank-sum test and two-tailed *p* values were used for statistical analysis. Asterisks compare respective *fzo-1(tm1133)* and wild-type N2 samples. * *p*<0.05; ** *p*<0.01; *** *p*<0.001; **** *p*<0.0001; n.s., not significant. (**D**) Heatmap of the MAI-2/IF1 protein expression in N2 and *fzo-1(tm1133)* strains at 12h and 24h post UV exposure is shown. The color bar depicts log2 expression fold change values. Individual protein changes and respective statistics for **A-D** are shown in Table S5.

**Figure S6.**
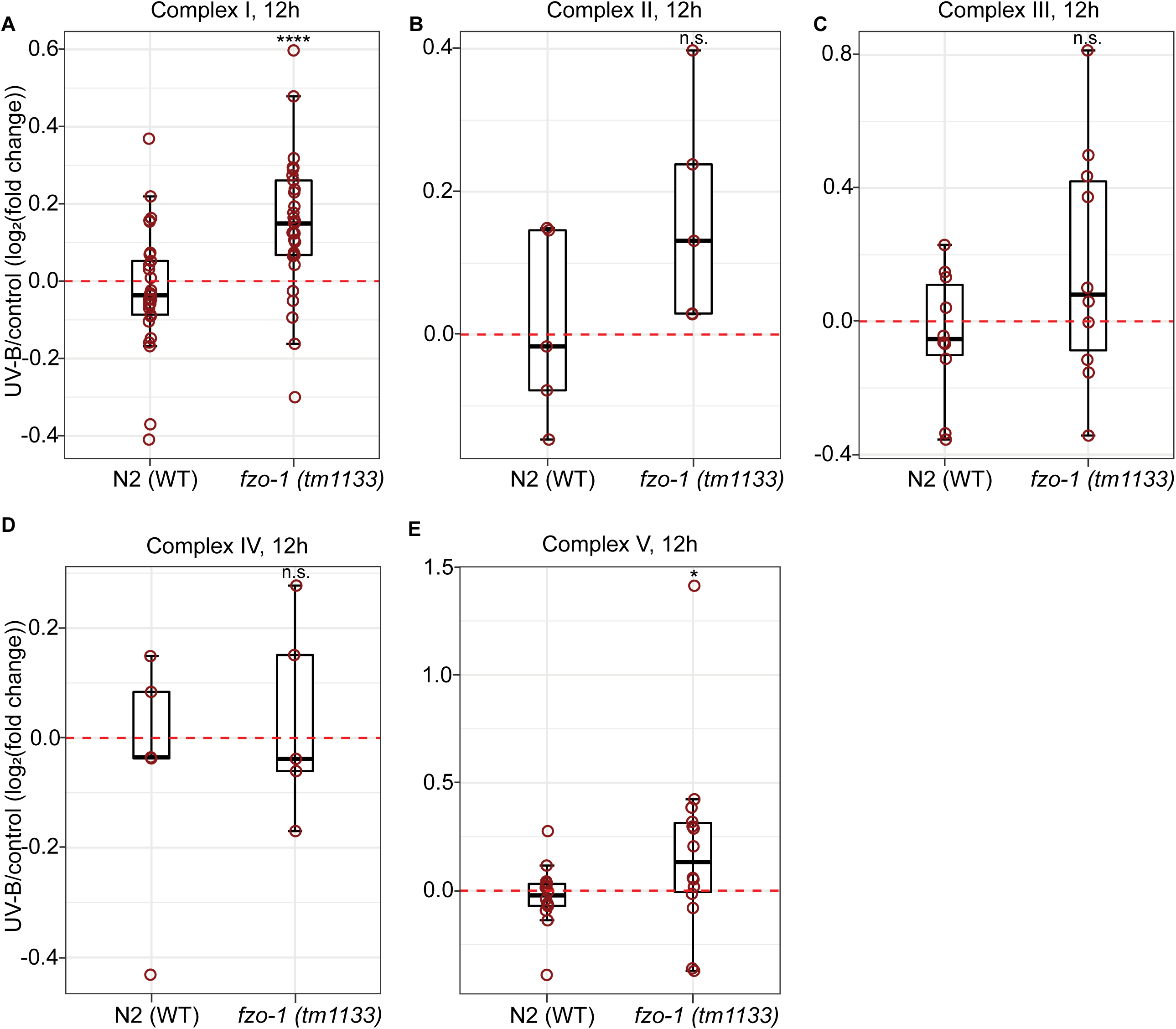
UVB triggers coordinated upregulation of ETC complexes I-V. UV treatment and proteomics sample collection was carried out as described in Figure 3. Box plots showing average Log2 expression fold changes of selected components of (**A**) Complex I (**B**) Complex II (**C**) Complex III (**D**) Complex IV and (**E**) Complex V at 12h post exposure to 300mJ/cm^2^ UV-B. In all cases, fold changes were calculated between UV-treated and respective time point matched controls samples. Red circles represent individual proteins and box plot parameters are as described in Figure 3 (A-C). Four independent pools of n=800 worms were analyzed for each condition. Wilcoxon rank-sum test and two-tailed *p* values were used for statistical analysis. Asterisks compare respective *fzo-1(tm1133)* and wild-type N2 samples. * *p*<0.05; ** *p*<0.01; *** *p*<0.001; **** *p*<0.0001; n.s., not significant. Individual protein changes and respective statistics for **A-E** are shown in Table S6.

**Figure S7.**
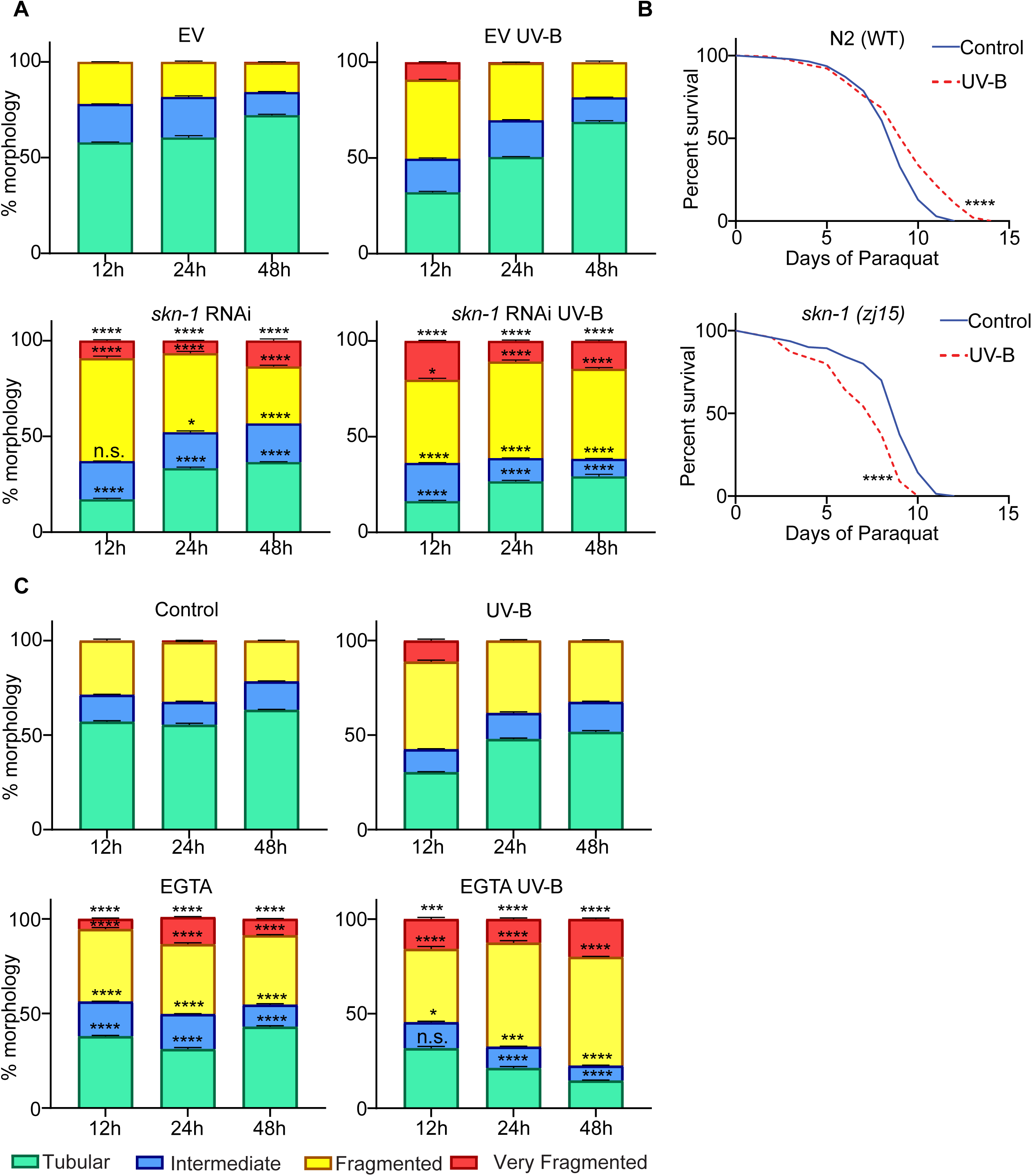
UVB-induced adaptive benefits require mitochondrial biogenesis and Ca^2+^ signaling. (**A**) Transgenic animals expressing GFP-tagged mitochondria in the body wall muscle (*myo-3p::gfpmit*) were grown on EV and *skn-1* RNAi from L1 stage and exposed to 850mJ/cm^2^ UVB on adulthood day 1 (AD1). Mitochondrial morphology was scored at 12h, 24h and 48h post UV-B treatment. Two-tailed unpaired t-test (with Welch’s correction) was used for the statistics. n=60 in each condition, mean and S.E.M values are presented. The asterisks compare respective morphologies of *skn-1* RNAi nematodes with time point- and treatment-matched EV control. (**B**) Wild-type (N2 Bristol strain) and *skn- 1(zj15)* mutant animals were pre-treated with 850mJ/cm^2^ UV-B at L4 stage and after 48h transferred to 5mM Paraquat (Sigma-856177) plates; survival was scored daily. Significance was measured by the Mantel-Cox test, and two-tailed *p* values were computed. Each group consisted of n=140 worms. (**C**) (*myo-3p::gfpmit*) transgenic animals were treated with UV-B (850mJ/cm^2^) on AD1 and immediately picked onto plates containing 50mM EGTA. The % of tubular, intermediate, fragmented, and very fragmented mitochondria were scored after 12h, 24h and 48h. Significance was measured by a two-tailed unpaired t-test (with Welch’s correction). n=60 in each condition, mean and S.E.M values are presented. The asterisks compare respective morphologies of EGTA plus nematodes with time point- and treatment-matched EGTA minus control. In **A-C** representative results of at least three independent experiments are shown.* *p*<0.05; ** *p*<0.01; *** *p*<0.001; **** *p*<0.0001; n.s., not significant.

**Figure S8.**
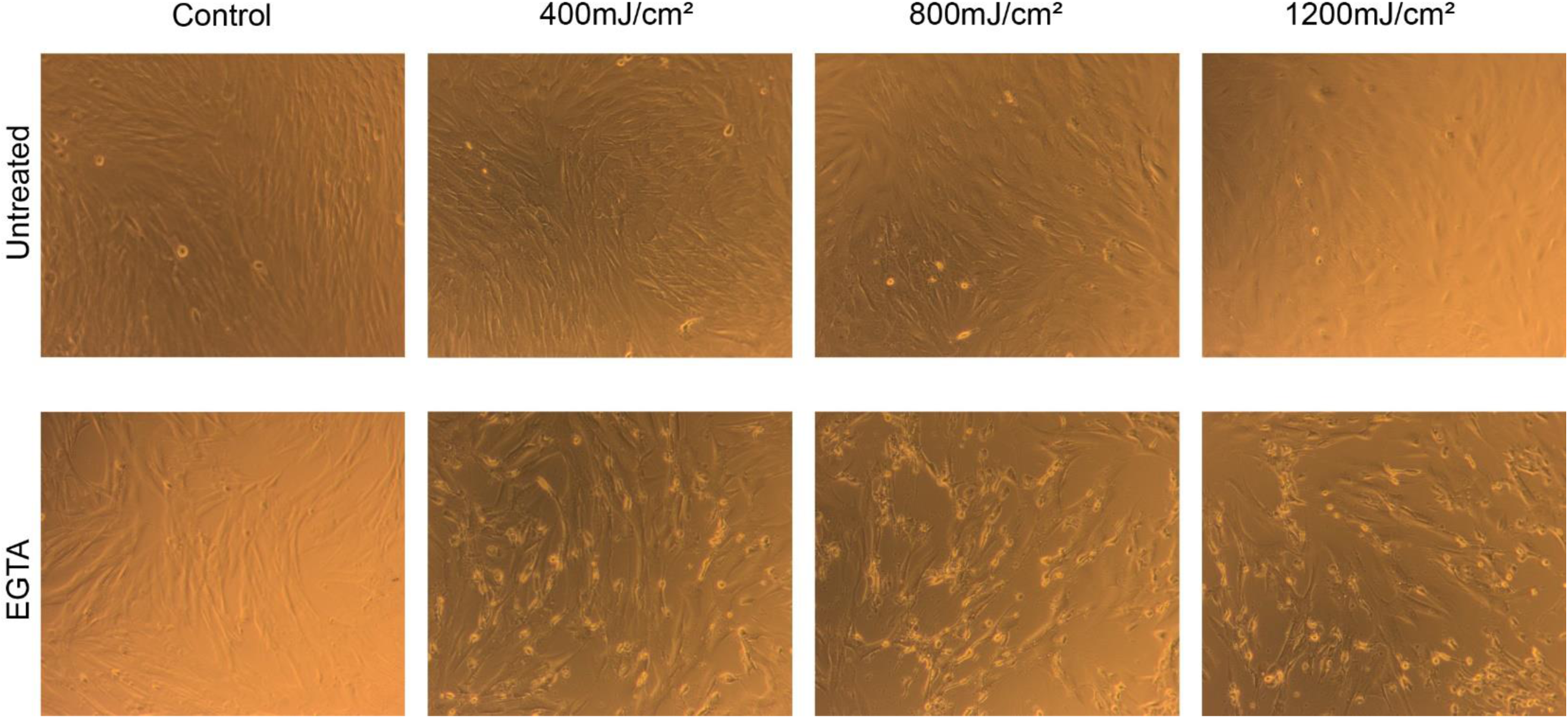
Ca^2+^ depletion sensitizes human skin fibroblasts to UVB toxicity. Representative microscopy images of human skin fibroblasts 24h post-treatment with 400 mJ/cm^2^, 800 mJ/cm^2^ and 1200 mJ/cm^2^ of UV-B and incubation in presence or absence of 2mM EGTA.

**Figure S9.**
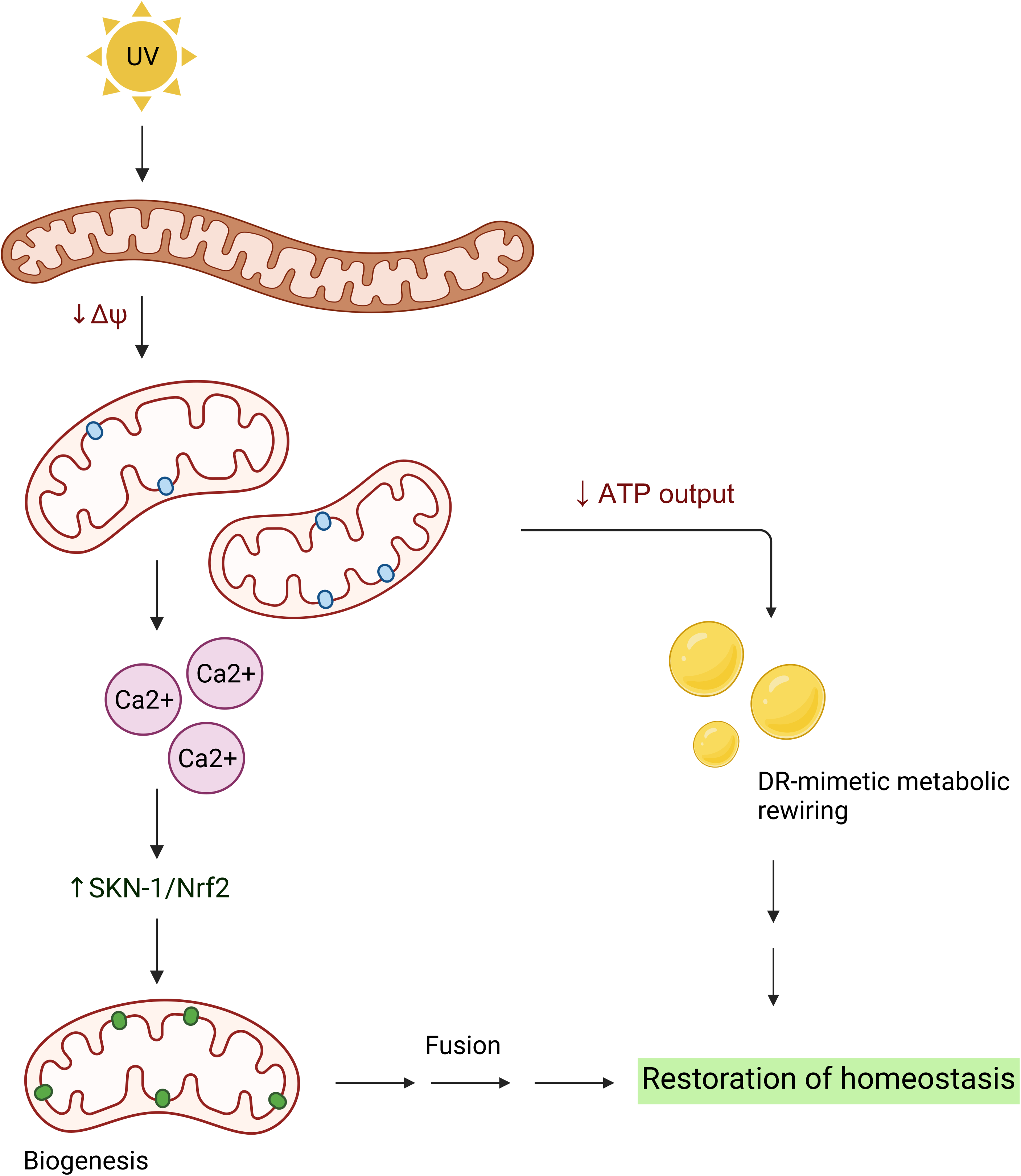
A model of UVB-induced metabolic rewiring response. UV-B light triggers mitochondrial network fragmentation and Ca^2+^ release via disruption of OXPHOS and distortion of mitochondrial ΔΨ. Ca^2+^ activates mitochondrial biogenesis via SKN-1/Nrf2, and newly generated ETC-rich mitochondria are integrated into the network by fusion to restore healthy homeostasis without lasting mitochondrial damage. While mitochondrial recovery takes place, the transient decline in ATP output triggers the DR-mimetic metabolic rewiring response, which warrants therapeutic exploration. Created with BioRender (biorender.com).

**Figure S10.**
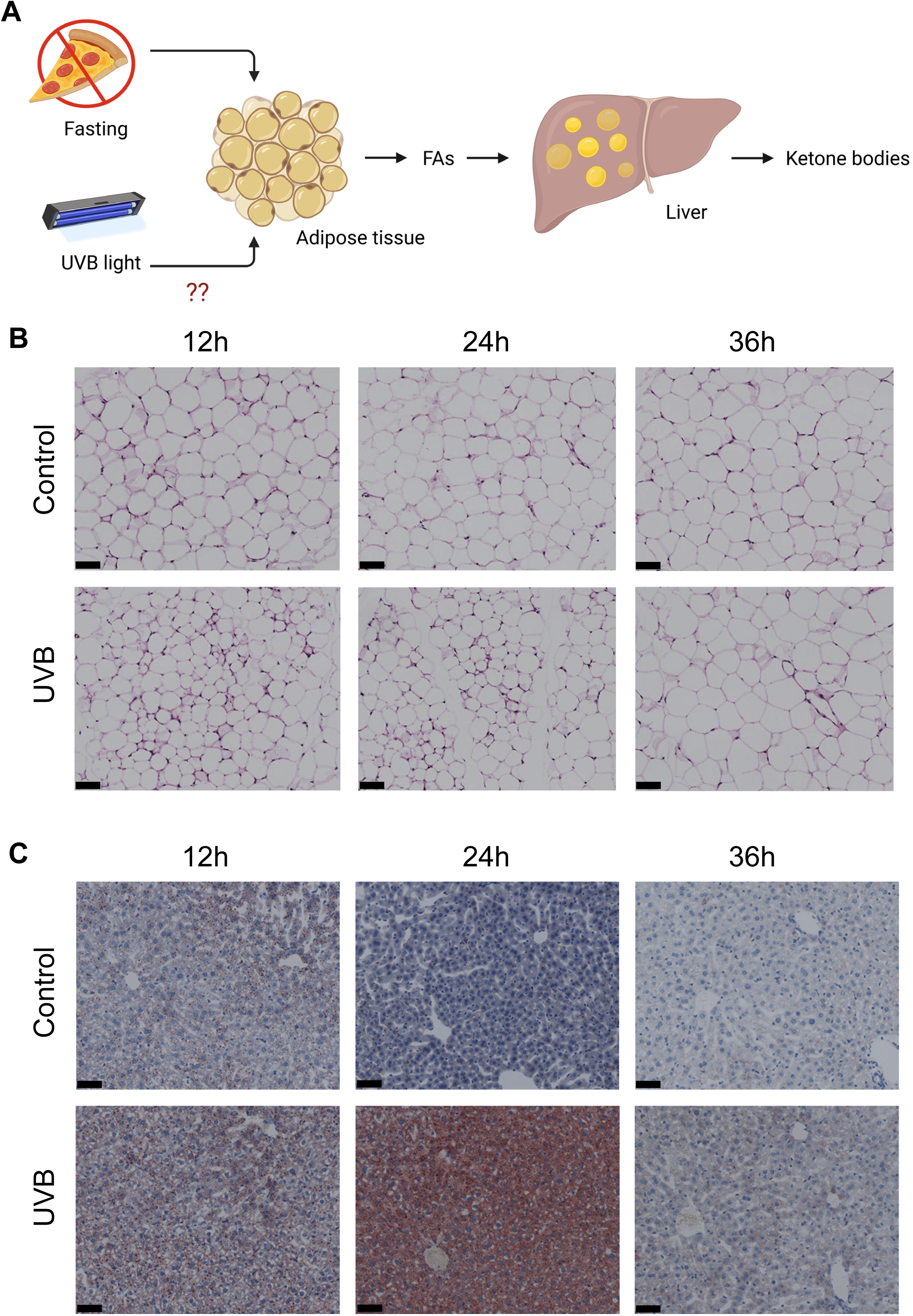
UVB irradiation of the skin triggers a systemic fasting-like effect in young mice. (**A**) Effects of fasting in the liver and white adipose tissue (WAT) are shown: in response to fasting, fatty acids are released from WAT, leading to shrinkage of WAT cells. These WAT-derived fatty acids are taken up by the liver where generation of ketone bodies takes place. (**B**) Hair was cleared from approx. 1/3 of total skin surface of young mice (back area, n=5 per condition), followed by irradiation with 100 mJ/cm^2^ of UVB. Control mice were treated similarly excluding UVB irradiation. White adipose tissue was collected at indicated times, and H&E staining results are presented showing reduced cell size in UVB-treated animals at 12h and 24h. (**C**) Oil Red O staining of liver tissue of mice described in (B). Transient increase of lipid levels is observed at 12h and especially at 24h post UVB exposure. Scale bar 50µm.

## Materials and methods

### *C. elegans* strains and culture conditions

The nematode strains used were HBR4 *goels3[myo-3p::SL1::GCamP3.35::SL2::unc543 ’UTR +unc-119(+)]*, N2 Bristol (wild type), SJ4100 *zcls13[hsp6p-GFP]*, AY131 *zcIs4[hsp- 4::GFP]*, SJ4103 *zcls-14[myo3p::GFP(mit)]*, CU5991 *fzo-1(tm1133)*, CU6372 *drp-1(tm1108)*, BR4006 *pink-1(tm1779)* and QV225 *skn-1(zj15)*. Strains were cultured under standard conditions^1,2^. Briefly, worms were grown on nematode growth media (NGM) with OP50 *E.coli* strain as a food source and at 20°C, unless stated otherwise. All nematode strains and the OP50 *E. coli* stock were obtained from the NIH funded Caenorhabditis Genetics Center (CGC).

### Exposure to ultraviolet light (UV) and ionizing radiation (IR)

For UV-B exposure, Medilux lamp UV 436 with 311nm wavelength was used and irradiation doses were monitored in real time by using Waldmann (Variocontrol) UV meter. The IR treatment was performed using gamma cell 40 (GC40 - Best Theratronics) irradiation chamber. The irradiation time was calculated based on the device-specific γ- radiation dose rate of approximately 0.8Gy per minute.

### Physiological stress treatments

For ER and oxidative stress treatments, the synchronized population of L1 animals was obtained by bleaching, and for heat stress tests – by filtering through the 11μM pore filters to minimize the stress prior to the testing. For heat stress exposure, the L1 worms were grown on 10cm NGM plates containing OP50 *E. coli* until L4 stage, treated with UV and IR and transferred to 35mm NGM plates, 3 plates per condition, 35 worms per plate. After 48h at 20°C the pates were transferred (in carton boxes) into an incubator running at 35°C. Plates were taken out and quickly scored for survival at indicated times followed by transfer back in the incubator. For ER and oxidative stress treatments, the UV- and IR-exposed L4 animals were transferred to fresh NGM plates containing OP50 *E. coli* and let recover for 48h at 20°C followed by transfer to 65mm NGM dishes containing 5mM Methyl viologen dichloride hydrate (Paraquat, oxidative stress) (Sigma-75365-73-0) or 10mM 1,4-Dithiothreitol (DTT, ER stress) (Sigma-3483-12-3). Two plates with 70 worms/plate were used for each condition. Survival was scored daily, and every three days remaining animals were transferred to fresh control, Paraquat or DTT dishes. For the young vs old comparison, animals were UV-treated on days 1 and 10 of adulthood, following the above procedure thereafter.

### Whole animal ATP measurements

ATP levels were measured as previously described^1^ with minor modifications. Briefly, worms were treated with UV-B and IR at young (non-gravid) adult stage with 5 plates of 50 worms used per condition. Worms were collected at indicated times, washed with S-basal, snap frozen in liquid nitrogen and stored at -80°C. Afterwards, sample preparation and ATP quantification assay was conducted with Roche ATP Bioluminescence Assay Kit HSII (Sigma) as indicated in the kit protocol and using Mithras LB940 plate reader (Berthold Technologies) for luminescence measurement. Three independent measurements of protein content for each sample were performed by using Nanodrop (ThermoFisher) device.

### Whole animal lipid staining

The staining of neutral lipids via Oil Red O (ORO) was performed as previously described^1^ with minor modifications. Briefly, adulthood day 1 worms on 10cm NGM plates were exposed to UV-B and IR and transferred to fresh 65mm NGM dishes containing OP50 *E. coli* at density of 30 animals/plate. For ORO protocol, the worms were collected at indicated times, washed in PBS and processed as previously described. The recording of the ORO signal was performs with Axio Scan.Z1 (Carl Zeiss) using 20x magnification and the HV-F202SCL color camera (Hitachi), and images were processed by Zen3.1 software (Carl Zeiss). Arbitrary units were set as follows and based on a previous report^3^: each worm raw image was greyscale-converted and later inverted for quantification. The worm was outlined and the mean grey value was measured using ‘’Analyse Measure’’ feature of IMAGE J (NIH) software. Each image background was measured four times, and later the average background value was subtracted from respective animal’s mean gray value. The arbitrary units (a.u.) plotting was done in GraphPad Prism 8 software.

### Mitochondrial morphology assessment

Assessment of mitochondrial morphology was performed as previously described^1^ with minor modifications. Briefly, filtered L1 stage *zcls-14[myo3p::GFP(mit)]* nematodes were grown on 10cm NGM dishes containing OP50 *E. coli* at a starting density of 500 animals per plate; on adulthood days 1 and 10 the animals were treated with UV and IR and picked on fresh 65mm NGM plates with *E.coli.* The worms were analyzed for % of tubular, intermediate, fragmented and very fragmented mitochondria at 12h, 24h and 48h post treatment. Each condition (time point) consisted of 3 independent replica plates containing 50 worms, and 20 worms per plate were examined.

### Proteomics analysis

Wild type and mitofusin mutant animals were treated with 300mJ/cm^2^ UV-B on adulthood day 1 and protein samples were collected after 12h and 24h. The dose of 300mJ/cm^2^ was determined experimentally as the maximal UV-B dose that did not lead to direct toxicity in high % of mitofusin mutants, which would otherwise hamper meaningful molecular analysis. 800 age-synchronized animals were used per replica and 4 independent replica samples were collected for each condition. Protein sample isolation, LC-MS/MS data collection in a data independent acquisition (DIA) mode and computational data analysis were performed as previously described^1^.

### Assessment of Ca^2+^ and UPR MT reporters

UPR MT *hsp-6p::GFP* reporter worms were age-synchronized, and L1 larvae were fed with HT115 *E. coli* containing empty vector (EV) or *mip-1* RNAi (known inducer of mitochondrial stress^1^). The worms were incubated at 20°C and treated with UV-B and IR as young (non-gravid) adults followed by microscopy analysis at 12h, 24h and 48h. For microscopy, nematodes were immobilized by cooling on ice for 20 minutes prior to imaging. Microscope and channels used were AxioZoomV16, brightfield TL and EGFP, 100x magnification was applied and Axiocam 503 with 15s exposure were used for image capturing. *myo-3p::SL1::GCamP3.35::SL2::unc543 ’UTR* Ca^2+^ reporter nematodes were grown on OP50 *E.coli* from L1 stage and until UV-B treatment at young (non-gravid) adult stage, and later (6h, 12h and 24h) assessed for GFP fluorescence with same microscope, channels and camera as *hsp-6p::GFP* worms with exposure of 20s. For both reporters, images were analyzed using IMAGE J software (NIH): whole animals were outlined and mean fluorescence was measured, followed by subtracting the average of four background measurements, and data analysis and plotting were done using Graph Pad Prism 8.

### Drug treatments in *C. elegans*

*myo3p::GFP(mit)* worms were age-synchronized and at L4 stage transferred to 60mm NGM plates seeded with OP50 *E.coli* and supplemented with 100μM and 200μM of rapamycin (AY-22989, LC Laboratories) at a density 50 worms per plate. After 24h, the worms were treated with UV-B and picked onto fresh rapamycin plates followed by mitochondrial morphology count at 12h, 24h and 48h post-treatment. Rapamycin was dissolved in DMSO and added directly to melted and cooled NGM agar with final DMSO concentration being 0.04%. An equal volume of DMSO was added to control agar plates.

Ethylene glycol tetraacetic acid (EGTA) (324626, Sigma) 0.5M concentrated stock was prepared. Next, 150 of 50mM EGTA dilutions were prepared for each plate. The liquid was gently swirled to cover the area of the plate and left to dry at RT. Due to anticipated minimal diffusion, the concertation was not adjusted by agar volume. The equal amount of EGTA free ddH2O was added on top of control plates. *E.coli* was seeded on dried plates and left at RT. *myo3p::GFP(mit)* worms were age-synchronized and grown under standard conditions until young (non-gravid) adult age, treated with UV-B, and transferred to 50mM EGTA plates followed by mitochondrial morphology scoring at 12h, 24h and 48h post-treatment.

### RNA interference (RNAi) treatments

RNAi exposure experiments were performed as previously described^1^. Briefly, HT115 *E. coli* bacteria containing gene-specific RNAi constructs were grown on double-resistant LB agar plates containing 0.2 mg/ml ampicillin and 0.01250 mg/ml tetracycline (K029.1and 0237.1, Roth). For treatment, overnight cultures from single colony were grown in LB medium with ampicillin. RNAi expression was induced by adding IPTG (16758, Sigma) to 2mM final concentration followed by 20 minute incubation at 37°C. Bacterial loans for worm feeding were then grown on NGM agar supplemented with 2.5mM IPTG and 0.2 mg/ml ampicillin.

### UV-B treatment in cell culture

BJ human skin fibroblasts (Coriell Institute) were maintained in DMEM high-glucose medium with L-glutamine and pyruvate supplemented with 10% FBS (all from Sigma). For UV-B exposure, cells were seeded in complete DMEM on flat-bottom 96-well plates at a density of 20,000 cells per well and avoiding rows on all edges of the plate due to light interference. Cells were let settle overnight followed by exposure to indicated doses of UV-B. All incubations and culturing were carried out in a humidified incubator (BBD6220, Thermo Scientific) at 37°C and 5% CO2. Primovert microscope and Axiocam ERc5s (both Carl Zeiss) were used for culture assessment and image capture at 10x magnification. For UV-B treatment, media was changed to 50 µl HBSS and after treatment, complete DMEM with indicated supplements (e.g. 2mM EGTA) was added.

### JC-1 mitochondrial membrane potential assay

The assay was adapted from the previous publication^1^. Briefly, 5μl of a filtered 55 µg/ml stock solution of the JC-1 dye (323590, Sigma) was added to each cell containing well to a final concentration of 5.5 µg/ml (8µM). Plates were wrapped in aluminium foil for light protection and incubated for 15 minutes at 37°C and 5% CO2. JC-1 medium was then removed, washed with 200µl DMEM, and replaced with fresh DMEM with phenol red. Fluorescence intensities at 530 nm (green, JC-1 monomers) and 590 nm (red, JC-1 aggregates) were measured by using a plate-reading spectrophotometer (Infinite M1000 Pro, Tecan).

### Proof-of-concept experiments in mice

#### Animals

Wild-type C57BL/6J mice (6–8 weeks old; 20.0 ± 2.0 g; female) were purchased from Charles River Laboratories (Beijing, China) and maintained under specific pathogen-free conditions. All animal procedures were approved by the Animal Ethical and Welfare Committee of Shenzhen University and conducted accordingly.

#### UV-B irradiation

Mice were anesthetized by isoflurane (2 % for induction and 1.2 % for maintenance; RWD, Shenzhen, China) in 1 L/min oxygen, and hair was removed by depilation from the back area. The UVB lamp (Telipu, Beijing Zhongyi Boteng Technology Company; intensity: 240±5 µW/cm2, wavelength: 308 nm) was placed 20 ± 1 cm above the skin surface. The UVB dose was measured by a photoelectric sensor (UV-B, Beijing Normal University Optoelectronics Technology Co., Ltd., China). The head of mice was covered to avoid eye damage. The irradiation was applied for 7 minutes reaching a total dose of 100 mJ/cm^2^, and control mice underwent same procedures excluding UVB exposure.

#### Tissue preparation and staining

Control and irradiated mice were sacrificed at three time points – 12h, 24h and 36h post-treatment, and liver and abdominal adipose tissue were isolated. For Hematoxylin/Eosin (H&E) staining, the adipose tissues were fixed with 4% paraformaldehyde (PFA), embedded in paraffin (JB-P5, Wuhan Junjie Electronics Co., Ltd., Wuhan, China) and sectioned at 4 µm using a microtome (RM2016, Leica, Shanghai, China). For Oil Red O staining, the liver tissues were embedded in OTC (Optimum Cutting Temperature Compound; SAKURA Tissue-Tek, USA) and sectioned at 7 µm using a cryostat (CRYOSTAR NX50, Thermo, USA). Images were captured with the Slide Scan System (SQS1000, TEKSQRAY, Shenzhen, China).

### Statistical analysis

The number of independent biological replicas for each experiment is stated in the figure legends, at least 3 replicas were included in all cases. Statistical analysis was performed with Prism 8 (Graphpad Software Inc.) or using custom designed R scripts (proteomics analysis). All statistics tests and values shown (e.g. mean ± S.E.M) are indicated in respective figure legends, significance was determined at *p*<0.05. All exact *p* values, *n* numbers and other statistics-relevant information are presented in the supplemented Source Data file.

## Bibliography

1. D’Orazio, J., Jarrett, S., Amaro-Ortiz, A. & Scott, T. UV radiation and the skin. Int J Mol Sci 14, 12222–12248 (2013).

2. Pfeifer, G.P. Mechanisms of UV-induced mutations and skin cancer. Genome Instab Dis 1, 99–113 (2020).

3. Slominski, A.T., Zmijewski, M.A., Plonka, P.M., Szaflarski, J.P. & Paus, R. How UV Light Touches the Brain and Endocrine System Through Skin, and Why. Endocrinology 159, 1992–2007 (2018).

4. Geldenhuys, S., et al. Ultraviolet radiation suppresses obesity and symptoms of metabolic syndrome independently of vitamin D in mice fed a high-fat diet. Diabetes 63, 3759–3769 (2014).

5. 5. Parikh, S., et al. Food-seeking behavior is triggered by skin ultraviolet exposure in males. Nat Metab (2022).

6. Ferguson, A.L., et al. Exposure to solar ultraviolet radiation limits diet-induced weight gain, increases liver triglycerides and prevents the early signs of cardiovascular disease in mice. Nutr Metab Cardiovasc Dis 29, 633–638 (2019).

7. Ermolaeva, M.A., et al. DNA damage in germ cells induces an innate immune response that triggers systemic stress resistance. Nature 501, 416–420 (2013).

8. Eckl, E.M., Ziegemann, O., Krumwiede, L., Fessler, E. & Jae, L.T. Sensing, signaling and surviving mitochondrial stress. Cell Mol Life Sci 78, 5925–5951 (2021).

9. Espada, L., et al. Loss of metabolic plasticity underlies metformin toxicity in aged Caenorhabditis elegans. Nat Metab 2, 1316–1331 (2020).

10. Rong, Z., et al. The Mitochondrial Response to DNA Damage. Front Cell Dev Biol 9, 669379 (2021).

11. Fang, E.F., et al. Nuclear DNA damage signalling to mitochondria in ageing. Nat Rev Mol Cell Biol 17, 308–321 (2016).

12. Hamsanathan, S., et al. Integrated -omics approach reveals persistent DNA damage rewires lipid metabolism and histone hyperacetylation via MYS-1/Tip60. Sci Adv 8, eabl6083 (2022).

13. Liu, Y.J., McIntyre, R.L., Janssens, G.E. & Houtkooper, R.H. Mitochondrial fission and fusion: A dynamic role in aging and potential target for age-related disease. Mech Ageing Dev 186, 111212 (2020).

14. Kozlowski, L., Garvis, S., Bedet, C. & Palladino, F. The Caenorhabditis elegans HP1 family protein HPL-2 maintains ER homeostasis through the UPR and hormesis. Proc Natl Acad Sci U S A 111, 5956–5961 (2014).

15. Weir, H.J., et al. Dietary Restriction and AMPK Increase Lifespan via Mitochondrial Network and Peroxisome Remodeling. Cell Metab 26, 884–896 e885 (2017).

16. Chaudhari, S.N. & Kipreos, E.T. Increased mitochondrial fusion allows the survival of older animals in diverse C. elegans longevity pathways. Nat Commun 8, 182 (2017).

17. Rambold, A.S., Cohen, S. & Lippincott-Schwartz, J. Fatty acid trafficking in starved cells: regulation by lipid droplet lipolysis, autophagy, and mitochondrial fusion dynamics. Dev Cell 32, 678–692 (2015).

18. Hsu, W.H., Lee, B.H. & Pan, T.M. Leptin-induced mitochondrial fusion mediates hepatic lipid accumulation. Int J Obes (Lond) 39, 1750–1756 (2015).

19. Han, S., et al. Mono-unsaturated fatty acids link H3K4me3 modifiers to C. elegans lifespan. Nature 544, 185–190 (2017).

20. Gartner, A., Milstein, S., Ahmed, S., Hodgkin, J. & Hengartner, M.O. A conserved checkpoint pathway mediates DNA damage--induced apoptosis and cell cycle arrest in C. elegans. Mol Cell 5, 435–443 (2000).

21. Lans, H., et al. Involvement of global genome repair, transcription coupled repair, and chromatin remodeling in UV DNA damage response changes during development. PLoS Genet 6, e1000941 (2010).

22. Lopes, A.F.C., et al. A C. elegans model for neurodegeneration in Cockayne syndrome. Nucleic Acids Res 48, 10973–10985 (2020).

23. Bennett, C.F., et al. Activation of the mitochondrial unfolded protein response does not predict longevity in Caenorhabditis elegans. Nat Commun 5, 3483 (2014).

24. Palikaras, K., Lionaki, E. & Tavernarakis, N. Coordination of mitophagy and mitochondrial biogenesis during ageing in C. elegans. Nature 521, 525–528 (2015).

25. Lapierre, L.R., et al. The TFEB orthologue HLH-30 regulates autophagy and modulates longevity in Caenorhabditis elegans. Nat Commun 4, 2267 (2013).

26. Mijaljica, D., Prescott, M. & Devenish, R.J. V-ATPase engagement in autophagic processes. Autophagy 7, 666–668 (2011).

27. Palikaras, K., Lionaki, E. & Tavernarakis, N. Coupling mitogenesis and mitophagy for longevity. Autophagy 11, 1428–1430 (2015).

28. Schwarz, J., Spies, J.P. & Bringmann, H. Reduced muscle contraction and a relaxed posture during sleep-like Lethargus. Worm 1, 12–14 (2012).

29. Rognoni, E., et al. Role of distinct fibroblast lineages and immune cells in dermal repair following UV radiation-induced tissue damage. Elife 10 (2021).

30. Dallam, R.D. & Hamilton, J.W. The Effect of Ultraviolet Irradiation on Mitochondrial Phosphorylation, Inorganic Phosphate-Atp Exchange and Atpase Activities. Radiat Res 22, 548–555 (1964).

31. Beyer, R.E. The effect of ultraviolet light on mitochondria. II. Restoration of oxidative phosphorylation with vitamin K1 after near-ultraviolet treatment. J Biol Chem 234, 688–692 (1959).

32. Miyazono, Y., et al. Uncoupled mitochondria quickly shorten along their long axis to form indented spheroids, instead of rings, in a fission-independent manner. Sci Rep 8, 350 (2018).

33. Zhao, Z., et al. Modulation of intracellular calcium waves and triggered activities by mitochondrial ca flux in mouse cardiomyocytes. PLoS One 8, e80574 (2013).

34. Beyer, R.E. The effect of ultraviolet light on mitochondria. IV. Inactivation and protection of the adenosine triphosphate-inorganic orthophosphate exchange reaction during far-ultraviolet irradiation. J Biol Chem 236, 3 (1961).

35. Ichikawa, N., et al. Activation of ATP hydrolysis by an uncoupler in mutant mitochondria lacking an intrinsic ATPase inhibitor in yeast. J Biol Chem 265, 6274–6278 (1990).

36. Chinopoulos, C., et al. Forward operation of adenine nucleotide translocase during F0F1-ATPase reversal: critical role of matrix substrate-level phosphorylation. FASEB J 24, 2405–2416 (2010).

37. Fernandez-Cardenas, L.P., et al. Caenorhabditis elegans ATPase inhibitor factor 1 (IF1) MAI-2 preserves the mitochondrial membrane potential (Deltapsim) and is important to induce germ cell apoptosis. PLoS One 12, e0181984 (2017).

38. Campanella, M., et al. Regulation of mitochondrial structure and function by the F1Fo-ATPase inhibitor protein, IF1. Cell Metab 8, 13–25 (2008).

39. Regmi, S.G., Rolland, S.G. & Conradt, B. Age-dependent changes in mitochondrial morphology and volume are not predictors of lifespan. Aging (Albany NY) 6, 118–130 (2014).

40. Amaro-Ortiz, A., Yan, B. & D’Orazio, J.A. Ultraviolet radiation, aging and the skin: prevention of damage by topical cAMP manipulation. Molecules 19, 6202–6219 (2014).

41. Tang, H.N., et al. Plasticity of adipose tissue in response to fasting and refeeding in male mice. Nutr Metab (Lond) 14, 3 (2017).

42. Zhang, X., et al. Fasting induces hepatic lipid accumulation by stimulating peroxisomal dicarboxylic acid oxidation. J Biol Chem 296, 100622 (2021).

43. Moller, L., Stodkilde-Jorgensen, H., Jensen, F.T. & Jorgensen, J.O. Fasting in healthy subjects is associated with intrahepatic accumulation of lipids as assessed by 1H-magnetic resonance spectroscopy. Clin Sci (Lond) 114, 547–552 (2008).

44. Kemeny, L., Varga, E. & Novak, Z. Advances in phototherapy for psoriasis and atopic dermatitis. Expert Rev Clin Immunol 15, 1205–1214 (2019).

45. Lee, T.L., et al. Vitamin D Attenuates Ischemia/Reperfusion-Induced Cardiac Injury by Reducing Mitochondrial Fission and Mitophagy. Front Pharmacol 11, 604700 (2020).

46. Ra, S.G., et al. Effects of Dietary Vitamin D Deficiency on Markers of Skeletal Muscle Mitochondrial Biogenesis and Dynamics. J Nutr Sci Vitaminol (Tokyo) 68, 243–249 (2022).

47. Wimalawansa, S.J. Vitamin D Deficiency: Effects on Oxidative Stress, Epigenetics, Gene Regulation, and Aging. Biology (Basel) 8 (2019).

48. Gorman, S., de Courten, B. & Lucas, R.M. Systematic Review of the Effects of Ultraviolet Radiation on Markers of Metabolic Dysfunction. Clin Biochem Rev 40, 147–162 (2019).

49. Singh, R.K., et al. The Patient’s Guide to Psoriasis Treatment. Part 1: UVB Phototherapy. Dermatol Ther (Heidelb) 6, 307–313 (2016).

50. Vangipuram, R. & Feldman, S.R. Ultraviolet phototherapy for cutaneous diseases: a concise review. Oral Dis 22, 253–259 (2016).

## Bibliography

1 Espada, L., et al. Loss of metabolic plasticity underlies metformin toxicity in aged Caenorhabditis elegans. Nat Metab 2, 1316–1331, doi:10.1038/s42255-020-00307-1 (2020).

2 Ermolaeva, M. A., et al. DNA damage in germ cells induces an innate immune response that triggers systemic stress resistance. Nature 501, 416–420, doi:10.1038/nature12452 (2013).

3 Han, S., et al. Mono-unsaturated fatty acids link H3K4me3 modifiers to C. elegans lifespan. Nature 544, 185–190, doi:10.1038/nature21686 (2017).

